# Predicting antimicrobial class specificity of small molecules using machine learning

**DOI:** 10.1101/2024.12.02.626313

**Authors:** Yojana Gadiya, Olga Genilloud, Ursula Bilitewski, Mark Brönstrup, Leonie von Berlin, Marie Attwood, Philip Gribbon, Andrea Zaliani

**Author notes:** **Corresponding author**: Yojana Gadiya, Fraunhofer ITMP.

## Abstract

Whilst the useful armory of antibiotic drugs is continually depleted due to the emergence of drug-resistant pathogens, the development of novel therapeutics has also slowed down. In the era of advanced computational methods, approaches like machine learning (ML) could be one potential solution to help reduce the high costs and complexity of antibiotic drug discovery and attract collaboration across organizations. In our work, we developed a large antimicrobial knowledge graph (AntiMicrobial-KG) as a repository for collecting and visualizing public *in-vitro* antibacterial assay. Utilizing this data, we build ML models to efficiently scan compound libraries to identify compounds with the potential to exhibit antimicrobial activity. Our strategy involved training seven classic ML models across six compound fingerprint representations, of which the Random Forest trained on the MHFP6 fingerprint outperformed, demonstrating an accuracy of 75.9% and Cohen’s Kappa score of 0.68. Finally, we illustrated the model’s applicability for predicting the antimicrobial properties of two small molecule screening libraries. Firstly, the EU-OpenScreen library was tested against a panel of Gram-positive, Gram-negative, and Fungal pathogens. Here, we unveiled that the model was able to correctly predict more than 30% of active compounds for Gram-positive, Gram-negative, and Fungal pathogens. Secondly, with the Enamine library, a commercially available HTS compound collection with claimed antibacterial properties, we predicted its antimicrobial activity and pathogen class specificity. These results may provide a means for accelerating research in AMR drug discovery efforts by carefully filtering out compounds from commercial libraries with lower chances of being active.

## 1. Introduction

Since their discovery, antibiotics have been primarily used to treat bacterial infections due to their ease of administration and potent antibacterial activity [1]. However, decades of liberal antibiotic use have led to a significant loss of effective treatment options. There is growing evidence that antimicrobial resistance (AMR) is an emerging threat to human health worldwide [2]. This has been highlighted in two comprehensive studies: the Antimicrobial Resistance report from 2016 [3] and the Global Burden of AMR study in 2019 [4, 5], among others [6]. To counter this, annual surveillance studies have generated and collated data, providing regional health institutions with opportunities to adapt and modify local prescribing trends and implement antimicrobial stewardship initiatives. Despite significant local, regional, and global efforts, the AMR burden remains at an all-time high. One major reason for the actual low rate of antibiotic authorization is the prolonged time required to develop drugs through the traditional drug discovery pipeline, coupled with the attrition of compounds that fail to reach the market [7]. Machine learning (ML), which can enable time-effective and efficient decision-making when presented with vast amounts of data, has the potential to improve drug discovery.

There is a pressing need to understand bacterial resistance phenotypes to develop effective AMR drugs. This necessity has driven efforts to manage AMR infections in both clinical and community settings. In clinical environments, omics experiments like whole-genome sequencing for antimicrobial susceptibility testing (WGS-AST) have shown the potential to provide rapid, consistent, and accurate predictions of known resistance phenotypes while offering rich surveillance data [8–11]. Meanwhile, judicious and controlled usage of existing medications in the community has also increased to combat this issue [12–14]. Understanding bacterial resistance mechanisms and characterizing the intrinsic pharmacokinetics and pharmacodynamics (PK/PD) features of drugs is essential in antibiotic drug discovery. Optimizing aspects such as drug metabolic stability, systemic half-life, and bioavailability often provides dosage advantages, enhancing the safety profile and minimizing the risks of rapid resistance emergence [15–17]. Therefore, such optimization processes require a swift turnover of novel synthetic candidates, increasingly identified using assistance from ML algorithms and models combined with de novo design methods [18–21].

The application of ML in AMR drug discovery has shown promising results, advancing from pre-clinical to clinical research stages [22]. Researchers have utilized ML algorithms to discover novel synergistic drug interactions from millions of potential combinations, thereby accelerating the development of combination therapies [23–26]. ML-informed computational approaches such as docking have been used to identify the activity of potential antibacterials against known microbial targets, as demonstrated by Chio *et al.* [27] and Alves *et al*. [28]. Similarly, predicting the activity of antimicrobial peptides against AlphaFold-predicted structures of microbial targets has been explored by Karnati *et al*. [29], among others [30, 31]. Susceptibility-related data is another source of data utilized by ML models to aid in selecting appropriate antibiotic therapy regimens for patients in clinical settings [32, 33]. These models help optimize treatment decisions by analyzing patterns in microbial resistance and patient-specific factors. Together, these efforts represent a shift in drug discovery, offering new avenues for the development of effective antimicrobial agents to combat the growing threat of AMR. Furthermore, by training mathematical models on empirical datasets, ML algorithms can predict the antibacterial activity of new compounds when presented with previously unseen data [19, 34]. The availability of public datasets, advances in computer engineering, and the proliferation of free and open-source ML libraries have deeply impacted this approach. However, many ML-based approaches serve the limitation of being predominantly focused on the phenotypic effects of drugs on target organisms [22, 35] rather than detailed molecular descriptions. This has led to underutilization of the chemo-physical characteristics of compounds [36] and insufficient attention to structural features [37] and general pharmacophoric features [38]. Leveraging these aspects can provide a more comprehensive understanding of drug behavior and efficacy, enhancing the development of novel therapeutic agents.

Our work addresses the previously mentioned limitations of existing ML models for predicting the antimicrobial activity of compounds. As part of the IMI AMR Accelerator project, COMBINE (https://amr-accelerator.eu/project/combine/), we gathered data on small molecules and their minimal inhibitory concentration (MIC) values from publicly available resources, creating a comprehensive database, the AntiMicrobial-KG. In this study, our primary goal was to develop ML models for predicting the activity of small molecules in the antimicrobial field, focusing on advanced pre-clinical drug discovery. This manuscript highlights the value of the utilization of structural-based molecular features for training ML models. Inherently, the insights from the model for these structural features will assist in developing promising compounds. We also built models to predict the broad-spectrum activity (antibacterial or antifungal) of novel compounds, facilitating efficient screening of compound libraries for experimental validation. Thus, by systematically integrating high-quality data to build transparent ML models, we can effectively demonstrate their applicability in assisting with AMR drug discovery.

## 2. Results

This section begins by presenting the schema of the AntiMicrobial-KG generated. In subsection 2.2, we investigated the performance of the models across different training datasets (SMOTE vs. non-SMOTE) generated using six chemical fingerprinting approaches (ECFP8, RDKit, MACCS, MHFP6, ErG, and ChemPhys). These datasets were analyzed with six different models from PyCaret: Naive Bayes (NB), Logistic Regression (LR), LightGBM, Decision Tree (DT), Random Forest (RF), and XGBoost. From these models, we select the best-performing one and, in subsection 2.3, evaluate it on external datasets, including commercial libraries, to identify potential antibacterial chemicals. Finally, subsection 2.4 discusses the limitations of our employed strategy.

### 2.1. AntiMicrobial-KG as a data warehouse for MIC bioassay endpoints

We built the AntiMicrobial-KG on the property-graph-based schema that is compliant with Neo4J. This data structure enabled systematic organization and accessibility of data, enhancing its reusability and compliance with FAIR principles. The graph consists of three types of nodes: Chemicals, Bacteria, and Pathogen class. It also includes two types of relationships (or edges), namely ‘shows_activity_on’ (connecting chemicals to bacteria) and ‘is_a’ (connecting bacteria to pathogen class), as illustrated in **Figure 1A**. At present, the AntiMicrobial-KG incorporates 82,867 nodes, with 81,490 chemicals tested across 1,373 bacteria and 831,432 relationships, constructing a harmonious interaction network. The distribution of the nodes and relationships in the AntiMicrobial-KG are summarized in **Figure 1B** and **Figure 1C**, respectively. To enable efficient chemical substructure searches, metadata for chemical representation in the form of SMILES and InChIKey are stored in the KG. Additionally, chemical classification from NPClassifier into classes, pathways, and superclasses for each chemical can be searched within the database. This comprehensive structure supports sophisticated querying and analysis, facilitating deeper insights into antimicrobial resistance patterns.

**Figure 1:**
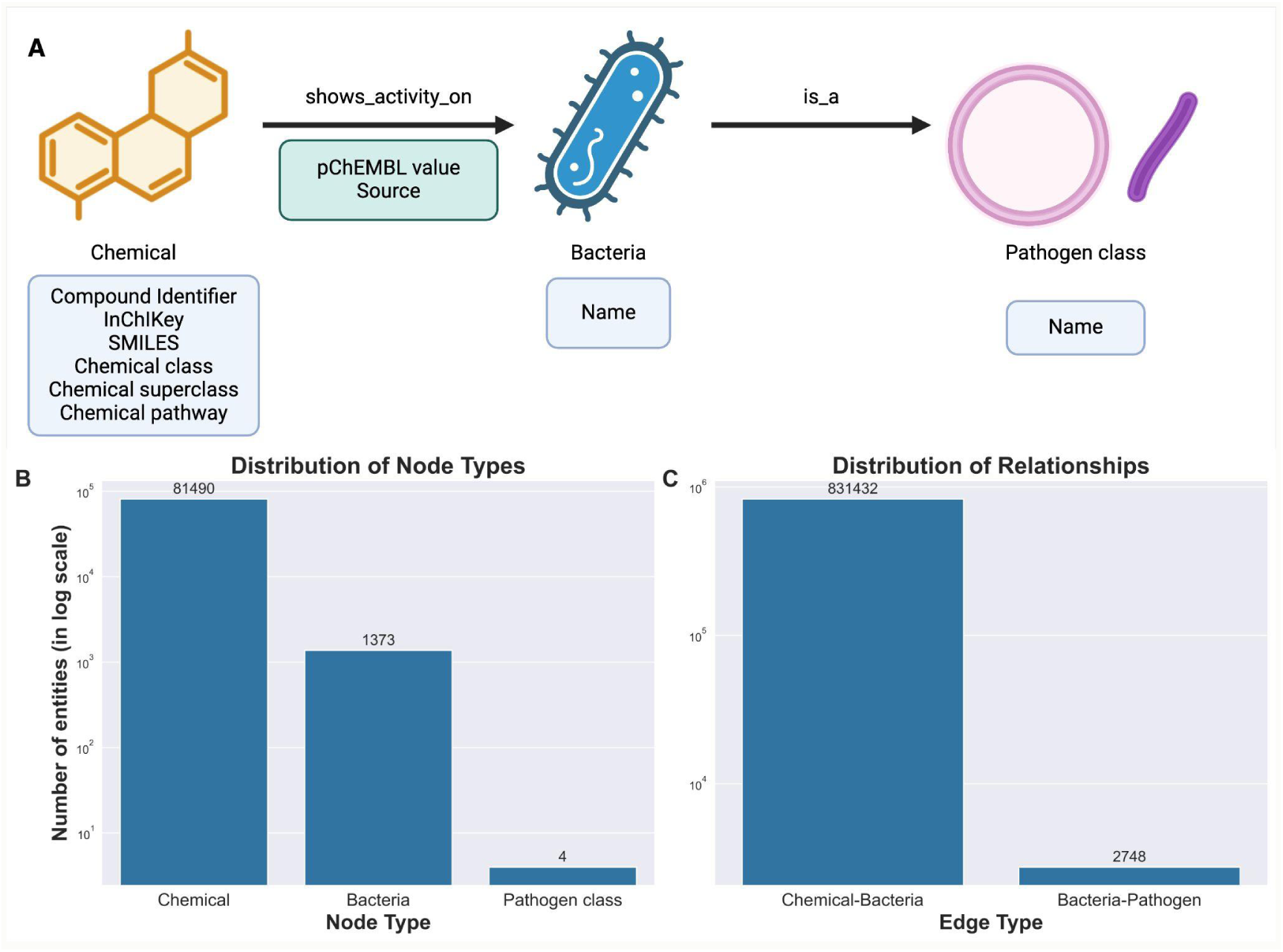
AntiMicrobial-KG schema and overview statistics. A) The schema of the AntiMicrobial-KG connecting the chemicals to bacteria and bacteria to pathogen class. Created with BioRender.com. B) The distribution of the number of nodes found in AntiMicrobial-KG. C) The distribution of edges in the AntiMicrobial-KG.

Next, we accessed the structural heterogeneity of the chemical space within the AntiMicrobial-KG by reducing the chemicals to their Murcko scaffolds. This investigation revealed a diverse chemical space within the AntiMicrobial-KG involving 24,506 unique Murcko scaffolds across 81,490 chemicals. We further classified the scaffolds based on their occurrence as ubiquitous and scarce within our dataset. As exemplified in **Figure 2**, Michael acceptor substructures were more commonly found in ubiquitous chemical scaffolds than in scarce ones (**Figure S1)**. Michael acceptors are compounds that contain an α,β-unsaturated carbonyl group, which can participate in Michael addition reactions. Molecular scaffolds containing chemically reactive groups such as pyridones, benzochromanones, and azetidinones target nucleophilic centers, often allowing for irreversible binding, a commonly observed property of natural products [39]. Moreover, Michael acceptor compounds are known to exhibit a variety of biological activities, including antimicrobial and antifungal properties, due to their ability to modify critical proteins and enzymes in microorganisms [40–42].

**Figure 2:**
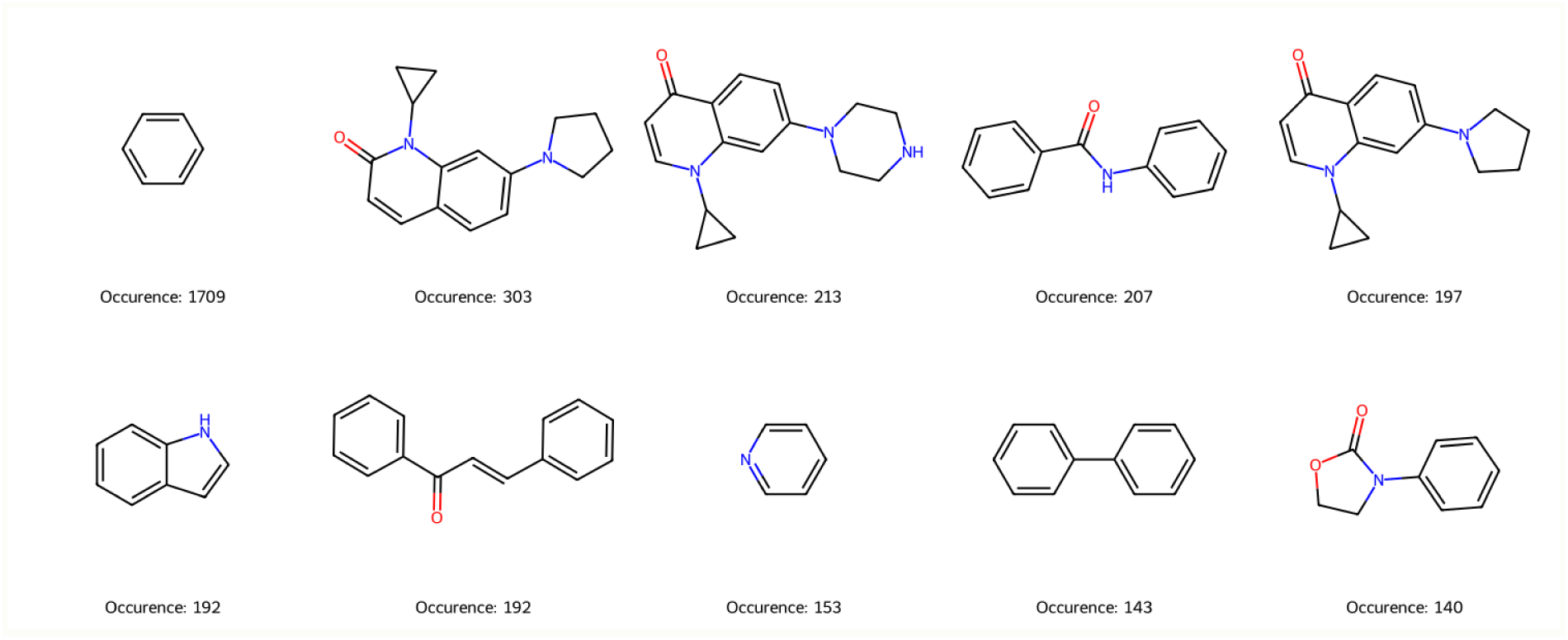
Top 10 ubiquitous generic Murko scaffolds identified in AntiMicrobial-KG. For each scaffold, the occurrence number in the AntiMicrobial-KG is depicted at the bottom.

Following the analysis of chemical diversity, we inspected the chemical space of the AntiMicrobial-KG by examining several aspects: Rule-of-Five (Ro5) compliance, the presence or absence of structural alerts, and the distribution of chemicals into classes, superclasses, and pathways across bacterial strains. We found that 75.2% (i.e., 55,814) of chemicals in the AntiMicrobial-KG violated at least one of the Ro5 guidelines, suggesting that a significant portion of the dataset consists of either natural products (NPs) originated chemicals or chemicals with poor oral bioavailability **(Table S1)**. To validate this, we assessed the NP-likeness of the chemicals and observed a substantial number of synthetic chemicals (25,313) compared to natural products (8,872), indicating that both NP origin and the poor bioavailability of synthetic compounds contribute to the Ro5 violations **(Figure S2)**. We also investigated the presence of structural alerts that include specific substructures known to be associated with toxicity or other undesirable properties. The overall ratio of chemicals with structural alerts to those without was approximately 7:3 across all pathogen classes. For chemicals with no structural alerts, the breakdown by pathogen class is as follows: 30.78% chemicals for Gram-positives, 27.93% for Gram-negatives, 31.39% for fungi, and 32.75% for acid-fast bacteria (**Table S1**). We further explored the characterization of chemicals within the AntiMicrobial-KG, grouping them into 411 chemical classes, 67 superclasses, and 7 biosynthetic pathways. This analysis revealed distinct patterns differentiating active from inactive chemicals across various pathogens, specifically at the chemical class level (**Figure 3**). For instance, 64 chemical classes, including tetracyclic and daphnane diterpenoids, steroidal alkaloids, and rotenoids, were exclusively found in Gram-positive active chemicals. Similarly, 34 classes, such as aeruginosins, pentacyclic guanidine alkaloids, and fasamycins and derivatives, were specific to Gram-negative active chemicals. For fungal pathogens, 42 chemical classes, including rhizoxins, ergostane steroids, and lipopeptides, were identified as active. Additionally, for acid-fast pathogens, 31 chemical classes, such as artemisinin, furans, polysaccharides, and tropane alkaloids, were uniquely associated with the active compounds. From a biosynthetic perspective, the polyketide pathway was the only pathway distinctly associated with Gram-positive active compounds compared to the inactive ones.

**Figure 3:**
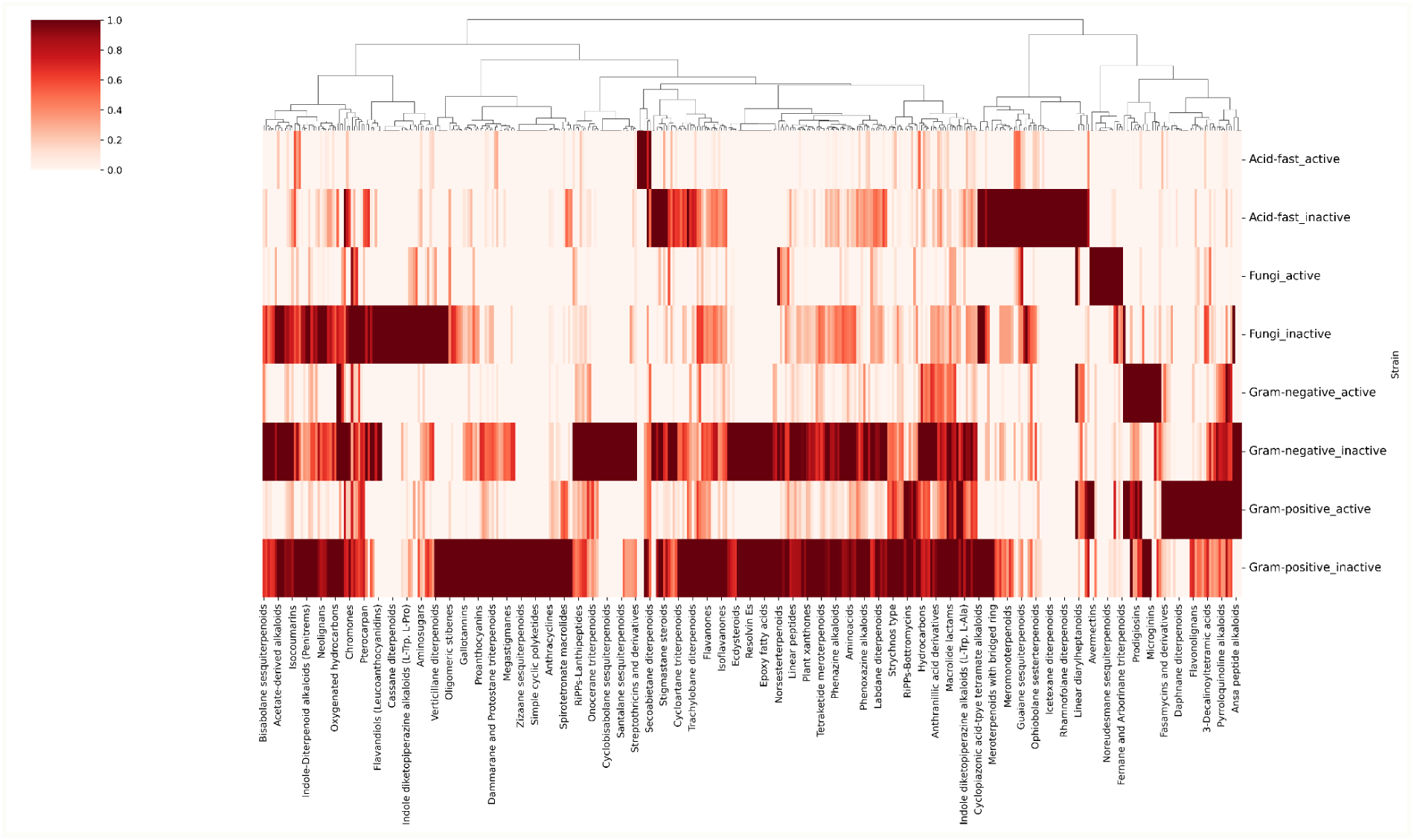
Cluster map of chemical classes across the four pathogen classes, each divided into two subclasses. From the cluster map, certain classes of chemicals are shown to be dominant (shown with dark red patches) across each of the pathogen categories, while others are underrepresented (shown with light red patches). Additionally, chemical class patterns are different between active and inactive chemicals for the same pathogen class.

### 2.2. MHFP6 with Random Forest outperforms other models

We began by developing a workflow to train and test a cohort of six (i.e. Naive Bayes, Logistic regression, Light Gradient Boosting Machine, Decision Tree, Random Forest and eXtreme Gradient Boosting) classical and explainable machine learning (ML) models. The implementation architecture encompasses data harmonization, model comparison, model selection, and optimization (see **Section 4.4** for more details). Our model comparison strategy allowed us to perform two independent assessments: first, to identify which model-fingerprint combination provided the best prediction results based on the Cohen Kappa score, and second, to evaluate the impact of implementing a SMOTE-based dataset on enhancing model prediction capabilities. Upon analyzing the model-fingerprint pair combinations, we found that, regardless of the fingerprinting approach used, the Random Forest and eXtreme Gradient Boosting (XGBoost) models consistently outperformed other models (**Figure 4**). In contrast, Naive Bayes and Logistic Regression models were the least effective for multi-class classification. Notably, there was a similar pattern in model outcomes (Cohen’s Kappa scores) between models using the RDKit fingerprint and those using MHFP6. Finally, models trained with ChemPhys and ErG fingerprints were the lowest performers, indicating that representing chemicals with fewer than 1,000 features (as these fingerprints do) is insufficient for capturing essential chemical characteristics compared to other fingerprints that utilize over 1,000 features. This insight underscores the importance of comprehensive feature representation in improving the predictive performance of ML models. The poor performance of the Naive Bayes models (Cohen’s Kappa score < 0.3) raises concerns about the implementation of the model in PyCaret and the predefined parameters used for training. The simplicity of the Naive Bayes model and its underlying assumptions about data distribution (e.g., Gaussian distribution for continuous features) and feature independence, which is rarely true in real-world data, further exacerbates its ineffectiveness in complex multi-class classification tasks. When comparing the SMOTE-trained models with classically trained models, we observed that the SMOTE-trained models exhibited an increase in performance, with an average improvement of 10% in Cohen’s Kappa score **(Table S2)**. To further evaluate the robustness of model predictions with SMOTE-based training, we trained the top two models (Random Forest and XGBoost) across all fingerprints using both the classic and SMOTE datasets **(Figure S3)**. This comparison consistently demonstrated that SMOTE training improved performance across various fingerprints, mirroring the initial observation.

**Figure 4:**
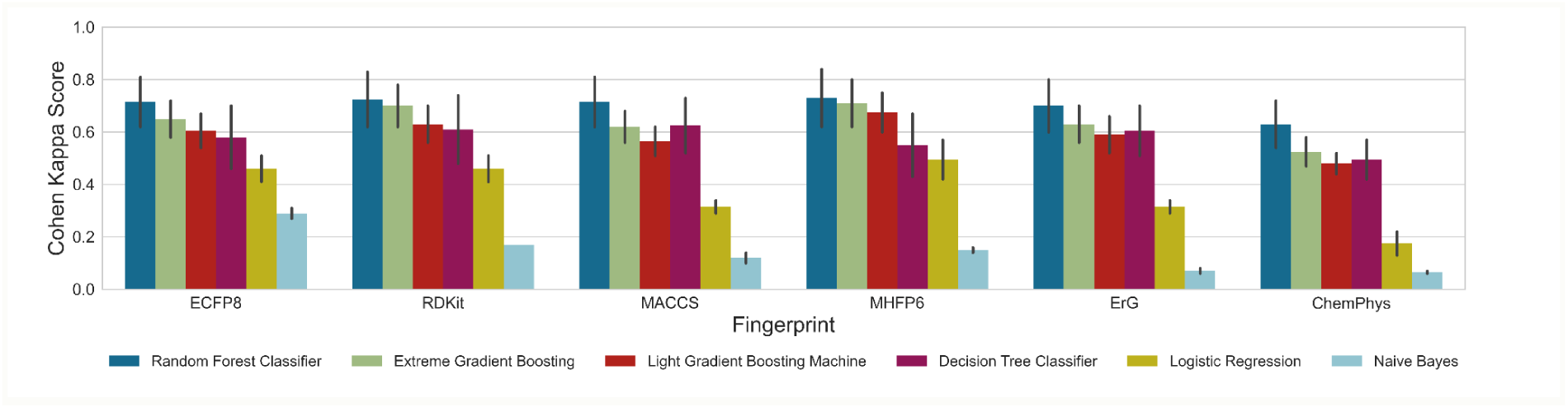
Comparison of the classic ML models across a spectrum of model-fingerprint combinations. The evaluation metric shown in this figure is based on the hold-out or validations set (20% of data), and models are trained on 60% of the data.

We identified Random Forest and XGboost as the top-performing models. These models were then subjected to hyperparameter optimization using SMOTE-trained data to enhance their predictive power. The specific hyperparameters optimized for training these two models are detailed in **Table S3.** The optimized models were then tested on the remaining 20% of test data. Among the two, Random Forest outperformed XGBoost, showing approximately 5% improvement across all metrics **(Table S4)**. **Table 1** highlights the performance of various fingerprints, with MHFP6 emerging as the best performer. This superior performance can be attributed to its unique chemical representation, which encodes the chemical using more than 2,000-bit vectors. This extensive encoding likely captures more detailed structural information, contributing to its enhanced predictive accuracy. In both models, the ChemPhys fingerprint showed the lowest performance, with 68.6% accuracy in random forests and 63.4% in XGBoost. This could be attributed to the lower number of features (i.e., 29 in ChemPhys vs 2048 in MHFP6) that correspond to the chemical. This small number might not be entirely sufficient to distinguish the activity of chemicals in final label classes (i.e. Gram-positive, Gram-negative, Acti-fast, Fungi, and Inactive).

**Table 1:** Random forest model performance on the test data. The table shows the average metric reported across all the five classes (Gram-positive, Gram-negative, Acid-fast, Fungi, and Inactive)

| Descriptors | Accuracy | Cohen Kappa | Macro precision | Macro recall | Macro F1 |
| --- | --- | --- | --- | --- | --- |
| MHFP6 | **0.759** | **0.678** | **0.775** | **0.756** | **0.764** |
| ECFP8 | 0.752 | 0.670 | 0.765 | 0.753 | 0.758 |
| RDKit | 0.743 | 0.659 | 0.752 | 0.746 | 0.749 |
| MACCS | 0.732 | 0.645 | 0.728 | 0.739 | 0.733 |
| ErG | 0.725 | 0.636 | 0.727 | 0.725 | 0.726 |
| ChemPhys | 0.686 | 0.585 | 0.671 | 0.680 | 0.675 |

All random forest trained fingerprint pairs demonstrated an average AUC-ROC score of 0.82. The MHFP6 fingerprint achieved the highest AUC-ROC score of 0.845, closely followed by the ECFP8-trained model with an AUC-ROC score of 0.843 **(Figure 5A)**. In contrast, the ChemPhys-trained model showed a lower AUC-ROC score of 0.79. For the best model pair, we further examined its ability to correctly classify chemicals into their respective pathogen classes using a confusion matrix **(Figure 5B)**. The true positive rate of Acid-fast pathogens was the highest at 0.81, while that of Gram-negative pathogens was 0.65. We also observed that overall, the model demonstrated a high precision or positive predictive value (PPV) and high negative predictive value (NPV) for all labels, indicating the model is robust and can correctly identify true positives and true negatives in the data (**Table S5**). Interestingly, 16% of Gram-negative active chemicals (i.e., 370 of 2,271 chemicals) were incorrectly classified as actives for the Gram-positive pathogen, and 5% of the Gram-positive active chemicals (i.e., 246 of 6,952 chemicals) were incorrectly classified as actives for Gram-negative pathogen. This misclassification can be attributed to two limitations of our current approach. According to our model, a chemical is favorable to be active only against a single pathogen class, which is disparate to real-world scenarios wherein a “broad spectrum” activity (i.e. activity across multiple pathogens) of chemicals is noticed. Secondly, the misclassification is due to the inability of the model to distinguish the MHFP6 fingerprint landscape of the two or more groups. One potential way to mitigate this issue is to increase the dataset used for training data, enabling the model to understand the MHFP6 chemical manifold space for better distinction.

**Figure 5:**
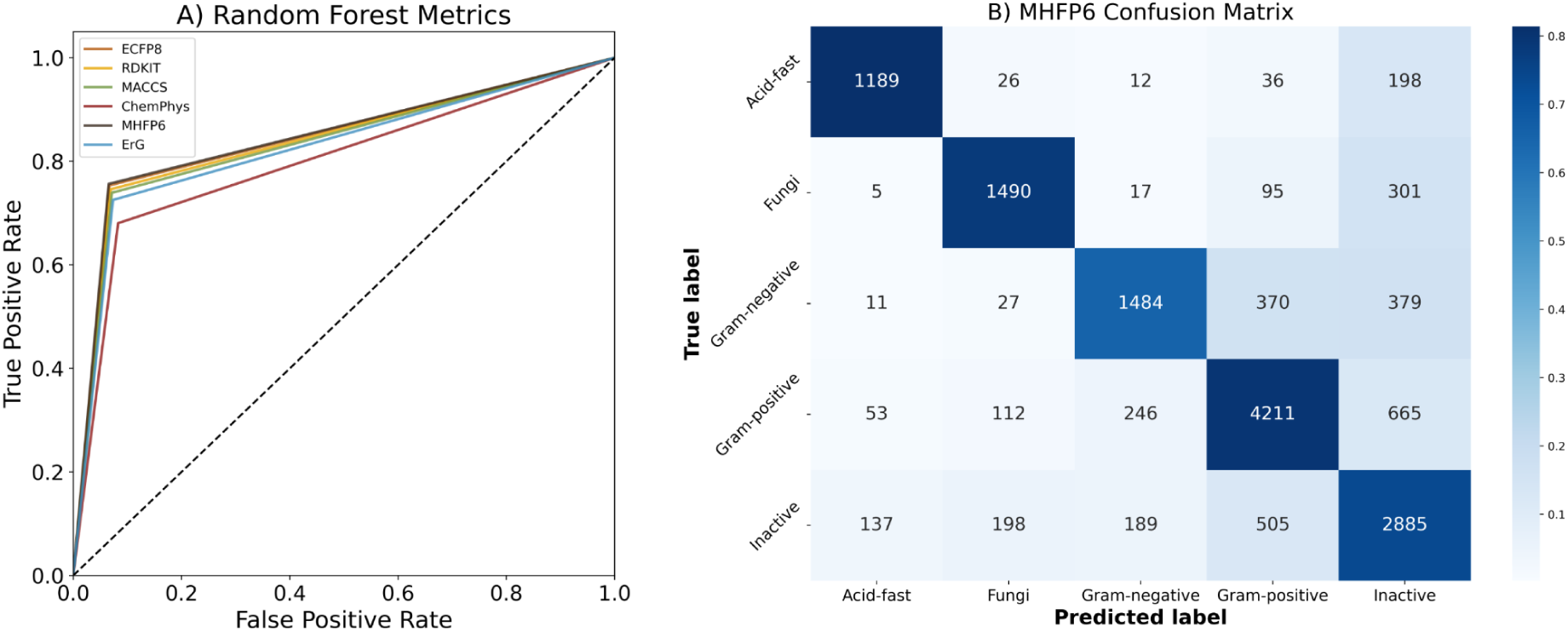
A) AUC-ROC sources of individual fingerprints trained with Random Forest. B) The confusion matrix of the best model is the MHFP6-trained Random Forest. The heatmap is color-coded based on the True Positive Rate (TPR) for each class, with annotations indicating the number of correctly classified chemicals.

Lastly, we inspected the feature importance of the model. Unfortunately, for most chemical fingerprints, the bit vectors are not easily translatable into chemical features that could directly inform future drug and lead optimizations (**Figure S4**). To address this, we leveraged the ChemPhys-trained model to correlate the target class with specific chemical properties, providing clues for improving antibacterial and antifungal drugs. Feature importance analysis from this model can be fundamental for interpreting the “trend rules” the model relies on to classify chemicals. These known “rules” can now be validated on a bigger chemical space than before and be used to generate novel hypotheses on important chemical properties. It is key to note that despite the ChemPhys-trained models demonstrating the lowest performance metrics overall, they still achieved an accuracy of 68.6%. Local label-specific feature importance analysis of this model revealed that chemical properties patterns allowed for clustering chemicals into three major buckets: Antibacterials (Gram-positive and Gram-negative pathogens), Anti-tuberculosis with acid-fast pathogens, and Antifungals with close patterns to those of inactive chemicals. Individual chemical properties such as hydrogen bond donors (HBD), the number of rings, LogP, and fraction SP3 played crucial roles in class predictability **(Figure 6)**. The heatmap allowed the identification of known and novel trend rules for antimicrobial activity. Firstly, SLogP, a critical parameter for solubility and cell permeability and a known factor influencing AMR activity [43], was identified by the model as a positive distinguishing feature between antibacterial and antifungal chemicals. Secondly, the presence and number of amide bonds, another well-established criterion for antibacterial activity particularly for Gram-negative strains [44, 45], was also identified and utilized by our model as a key distinguishing factor. Thirdly, the hydrogen bond donor (HBD) count (exact or Lipinski filtered) is more sensitive for Gram-type pathogens than for fungi or acid-fast. To the best of our knowledge, this specific aspect has not been highlighted in any previous antimicrobial activity prediction models. Another example of novel trends we identified was the role of aliphatic rings (carbocyclic or heterocyclic). The model used the number of these rings to differentiate Gram-positive active chemicals from Gram-negative active chemicals. Additionally, the number of saturated rings helped identify antifungal compounds, while the presence of saturated heterocycles was linked to anti-tuberculosis activity. Finally, a number of different descriptors dealing with structural complexity, polar surface, and atomic nature have been chosen to dissect inactive compounds.

**Figure 6:**
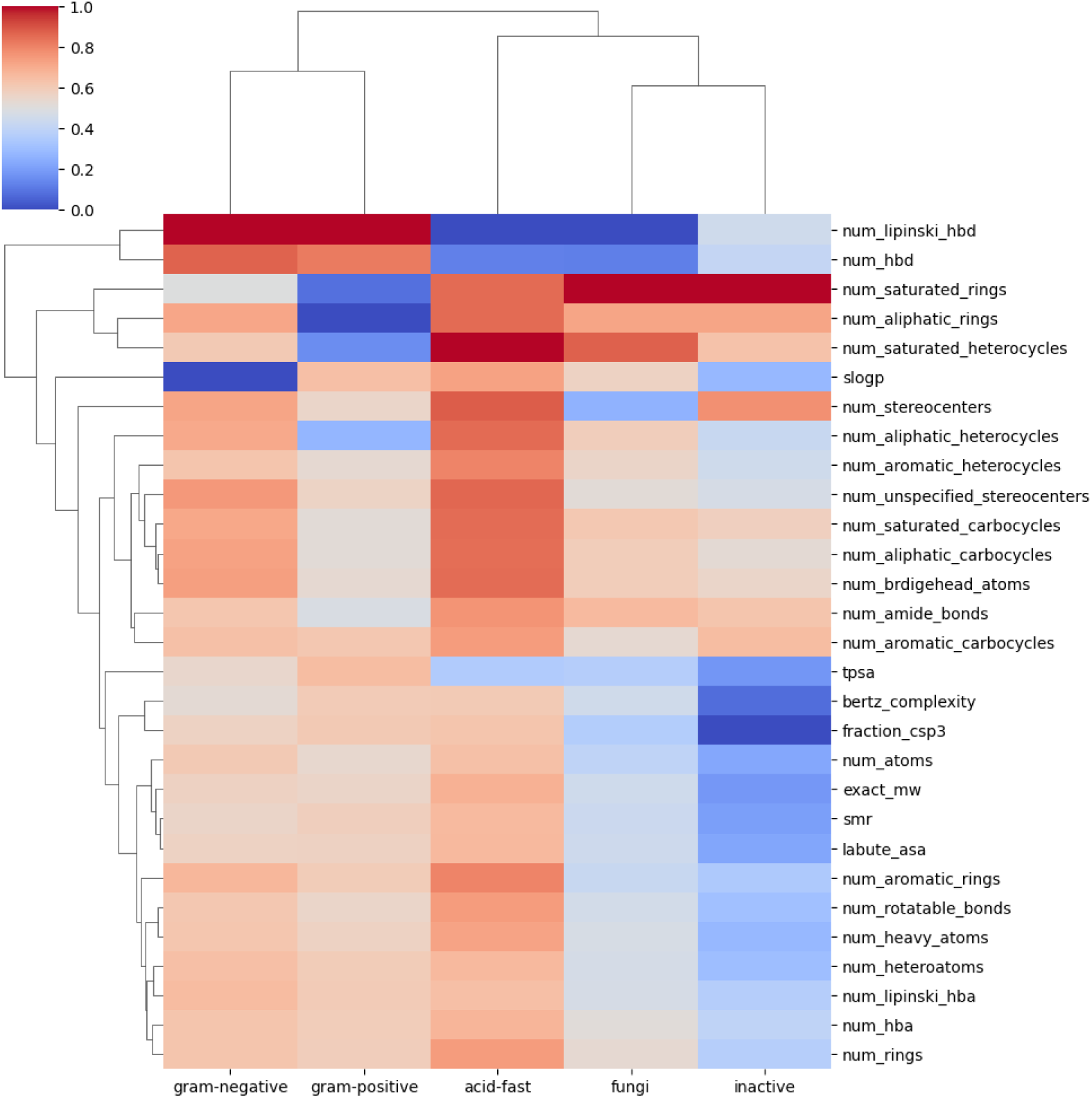
Heatmap of the ChemPhys importance across the five label classes. The physicochemical descriptors are ranked in a top-down manner based on their importance in describing the target or activity class of the dataset. The rows indicate the ChemPhys properties of the chemical, the columns indicate the activity class or label, and the color intensity indicates the importance of the normalized feature for each ChemPhys-activity pair.

### 2.3. Testing the model with external compound libraries

Finally, we tested the best-performing model, Random Forest trained with MHFP6 fingerprint, on commercial compound libraries to evaluate their *in-silico* antimicrobial activity. This testing phase was crucial for assessing the model’s practical relevance and potential effectiveness in identifying novel or repurposed compounds useful for combating AMR. Additionally, by applying the model to a diverse range of commercially available compounds, we aimed to determine ML model robustness and reliability in real-world scenarios.

The complete EU-OS library, comprising the ECBL and Bioactive sets, was screened against seven microbial pathogens: *Candida auris* DSM21092, *Staphylococcus aureus* ATCC 29213, *Pseudomonas aeruginosa* group, *Candida albicans* ATCC 64124, *Enterococcus faecalis* ATCC 29212, *Aspergillus fumigatus* ATCC 46645, and *Escherichia coli* ATCC 25922. To classify the compounds based on their activity, an arbitrary threshold of ≥ 50% inhibition (at 50 µM) was applied to distinguish active from inactive compounds. Using this criterion, over 95,000 compounds from the ECBL set were labelled as inactive, while approximately 1,000 compounds were classified as active **(Figure 7)**. Specifically, 983 compounds were found to be active against *Staphylococcus aureus*, 1 compound against *Pseudomonas aeruginosa*, 946 compounds against *Candida auris*, 34 compounds against *Enterococcus faecalis*, 198 compounds against *Aspergillus fumigatus*, 131 compounds against *Candida albicans*, and 26 compounds against *Escherichia coli*. From the pathogen class perspective, 99.7% of the compounds were inactive, and 0.3% of the compounds were active for Fungi. For Gram-negative pathogens, 99.9% of the compounds were inactive, and 0.01% of the compounds were active, while for Gram-positive pathogens, 98.9% of the compounds were inactive, and 1.1% of the compounds were active.

**Figure 7:**
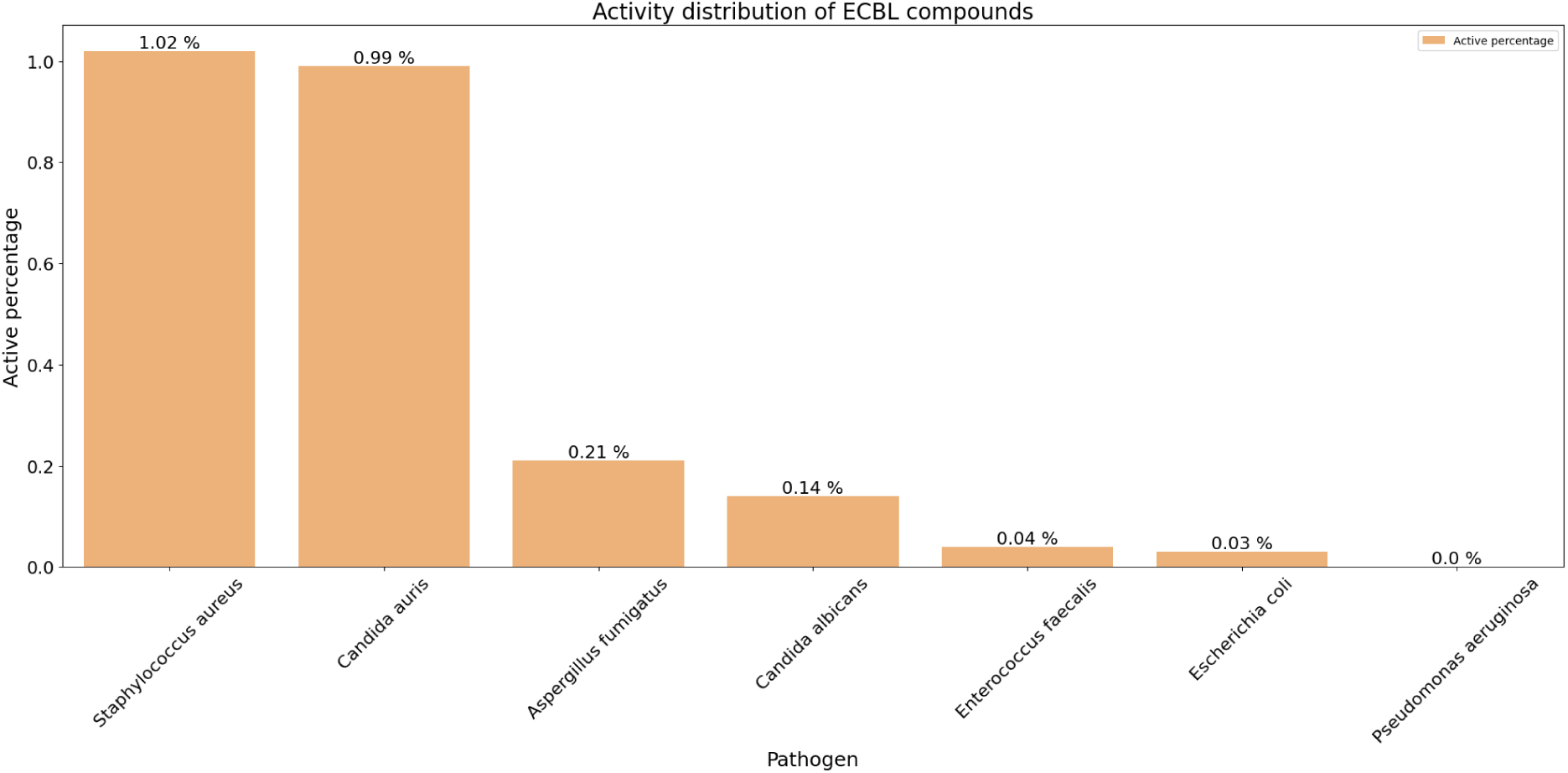
The percentage of experimental active compounds in the ECBL Library for each bacterial strain. The threshold was set to 50% inhibition with compound activity value > 50% classified as actives, and compounds with activity value <50% were classified as inactive.

Analogously, for the Bioactive set, approximately 4,700 compounds were classified as inactive, while the remaining 300 compounds were active **(Figure 8)**. Among these, 313 compounds were found to be active against *Staphylococcus aureus*, 13 compounds against *Pseudomonas aeruginosa*, 275 compounds against *Candida auris*, 123 compounds against *Enterococcus faecalis*, 145 compounds against *Aspergillus fumigatus*, 120 compounds against *Candida albicans*, and 52 compounds against *Escherichia coli*. From the pathogen class perspective, 95% of the compounds were inactive, and 5% of the compounds were active for Fungi. For Gram-negative pathogens, 98.9% of the compounds were inactive, and 1.1% of the compounds were active, while for Gram-positive pathogens, 94% of the compounds were inactive, and 6% of the compounds were active. Compared to the ECBL set, the Bioactive set exhibited a higher experimental hit rate. This could be attributed to several factors, such as the novelty of the ECBL compounds, which may not yet be optimized for cellular permeability. Additionally, the Bioactive set is inherently biased toward “activity,” even if on different protein targets, which could explain its higher hit rate. Another potential factor is the greater structural diversity present in the Bioactive set compared to the ECBL **(Figure S5)**. The ECBL, by design, aimed to sample a mini-family of 6–8 compounds around each collection component to ensure a “mini SAR” (structure-activity relationship) in case a hit was identified.

**Figure 8:**
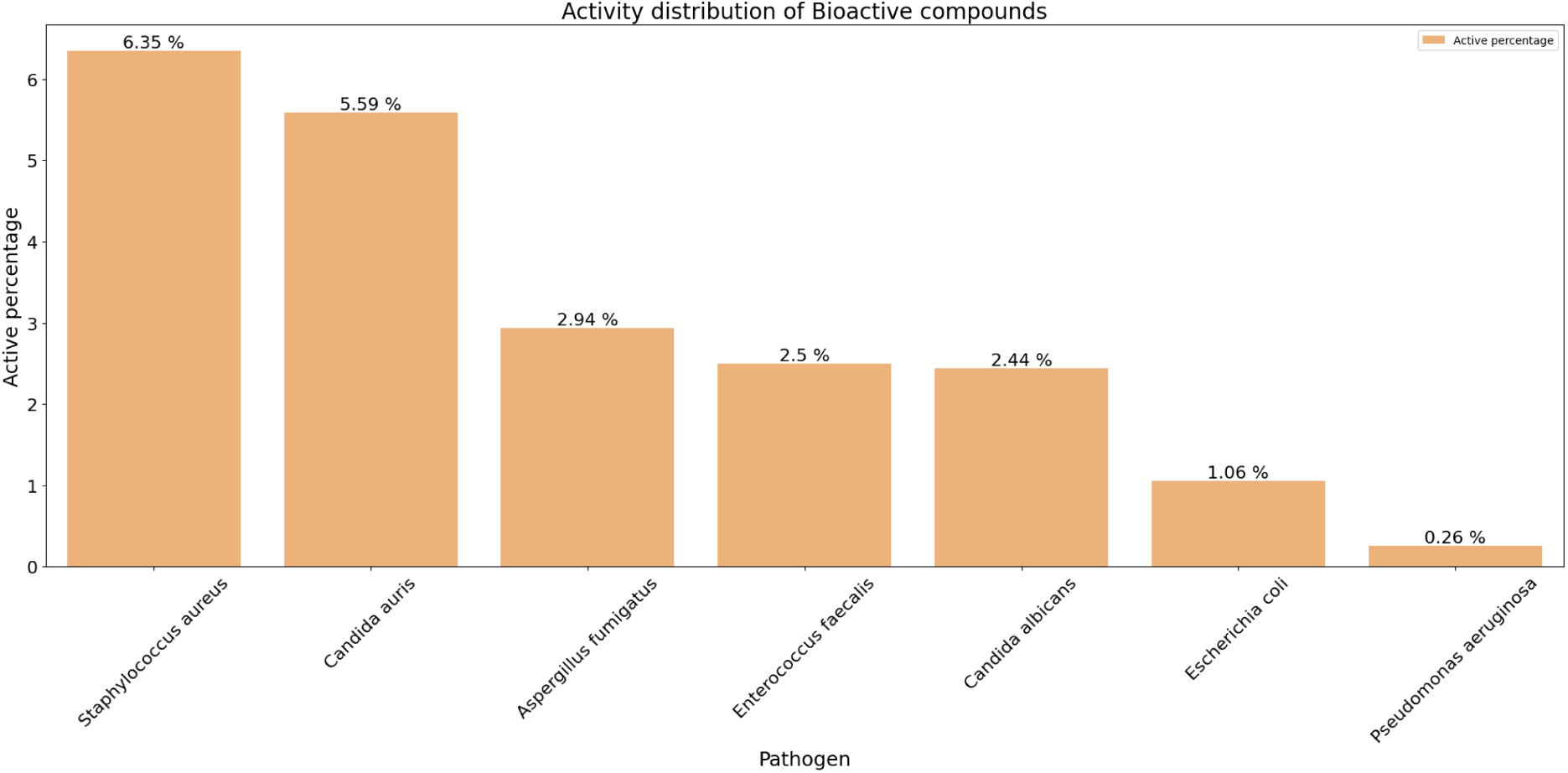
The percentage of experimental active compounds in the Bioactive Library across each bacterial strain. The threshold was set to 50% inhibition with compound activity value ≥ 50% classified as actives, and compounds with activity value <50% were classified as inactive.

Model predictions for the ECBL set indicated that 90% of the compounds were inactive, with 3.89% active against Gram-positive bacteria, 2.7% against acid-fast bacteria, 1.53% against fungi, and 0.99% against Gram-negative bacteria. In contrast, for the Bioactive set, 81.8% of the compounds were inactive, while 11% were active against Gram-positive bacteria, 3.13% against acid-fast bacteria, 1.99% against fungi, and 2.13% against Gram-negative bacteria. Next, we evaluated the hit rate of the model predictions in comparison to the experimental results, as summarized in **Table 2**. This analysis focused on examining the number of active compounds identified by both the model and the experiments. In most cases, the model’s hit rate exceeded that of the experimental results. Furthermore, a significant difference was observed when comparing the two libraries. For the ECBL library, the difference between the model-predicted hits and experimental hits was 0.17, whereas for the Bioactive library, this difference was markedly higher, at 18.37. The highest hit rate was observed for the Fungal strains, followed by the Gram-positive and Gram-negative, respectively. We also observed a similar trend in the hit rate difference for each pathogen class (**Table S6**). As with any ML model, the higher hit rate observed for the Bioactive library can be attributed to the similarity of its chemical space to the model’s training dataset.

**Table 2:** Comparison of the HitRate associated with the two library sets: ECBL and Bioactive. For each strain tested in EU-OS, the HitRate for the model predictions and experimental results are reported.

| Pathogen class | Experimental Strain | ECBL Model HitRate<br>(Experimental HitRate) | Bioactive Model HitRate<br>(Experimental HitRate) |
| --- | --- | --- | --- |
| Gram-positive | *Staphylococcus aureus* | 1.09% (1.02%) | 14.81% (6.35%) |
| Gram-positive | *Enterococcus faecalis* | 0.16% (0.04%) | 8.52% (2.49%) |
| Gram-negative | *Pseudomonas aeruginosa* | 0% (0.001%) | 5.71% (0.26%) |
| Gram-negative | *Escherichia coli* | 0.32% (0.03%) | 10.48% (1.06%) |
| Fungi | *Candida auris* | 1.29% (0.98%) | 42.86% (5.58%) |
| Fungi | *Aspergillus fumigatus* | 0.41% (0.21%) | 36.73% (2.94%) |
| Fungi | *Candida albicans* | 0.34% (0.14%) | 30.61% (2.43%) |

In addition to assessing the hit rate success of the model, we quantified its potential impact on future predictions, particularly in terms of cost savings. A key consideration in this analysis was addressing the question: How much money could be saved by leveraging our ML models? To explore this, we compared the compounds predicted to be active by the model against the entire compound library. In a typical scenario, such as with the EU-OS library, all compounds would need to be tested against all pathogens to identify a hit—defined as a compound exhibiting activity against a specific pathogen class or strain. However, by utilizing the model prior to experimental screening, a significantly smaller subset of compounds needs to be tested to achieve a comparable hit rate. For example, in the EU-OS library, the highest hit rate for *Staphylococcus aureus* was achieved with a library size of 100,000 compounds (**Figures 7** **and 8**). The model was able to identify nearly 25% of these active compounds by validating only 11% of the library (**Table 3**). On a broader scale, for Gram-positive pathogens, testing just 11% of the compound library was sufficient to capture over 30% of the hits, on average. Similarly, for Gram-negative pathogens and Fungi, as little as 2% of the compound library needed to be tested to identify active hits. This highlights the efficiency and cost-effectiveness of incorporating ML models into the screening process, significantly reducing the experimental workload while maintaining high hit rates.

**Table 3:** Impact quantification of the model predictions on the Bioactive library. For each strain tested in EU-OS, the number of actives found with a small percentage of screening library is shown.

| Pathogen class | Experimental Strain | True active compound percentage | Screening library percentage |
| --- | --- | --- | --- |
| Gram-positive | *Staphylococcus aureus* | 25.6% | 10.96% |
| Gram-positive | *Enterococcus faecalis* | 37.4% | 10.96% |
| Gram-negative | *Pseudomonas aeruginosa* | 46.1% | 2.13% |
| Gram-negative | *Escherichia coli* | 21.1% | 2.13% |
| Fungi | *Candida auris* | 15.3% | 1.98% |
| Fungi | *Aspergillus fumigatus* | 24.8% | 1.98% |
| Fungi | *Candida albicans* | 25% | 1.98% |

On the other hand, for the 32,000 compounds from Enamine, we found ∼90% of the compounds to demonstrate inactivity for antibacterial, antifungal, and antituberculosis activities. For the remaining 10%, many compounds are predicted to show activity against Gram-positive (∼68%, 2,276 compounds) and acid-fast (∼21%, 719 compounds) pathogens **(Figure 9)**. In addition to looking at prediction probabilities like that of EU-OS prediction analysis, we looked at the cosine similarity between the MHFP6 fingerprints of library compounds and the training dataset to assess the applicability domain of each predicted dataset. A median cosine similarity of 0.71 was reported for the compound library, with six compounds demonstrating a perfect cosine similarity of 1 to the training datasets. If we focus on high-confidence predictions (prediction probabilities > 0.5), we retained 27 compounds from the library with predicted activity against Gram-positive (8 of 33 compounds), Fungi (11 of 33 compounds), Gram-negative (1 of 33 compounds), and Acid-fast (7 of 33 compounds pathogens **(Table S7)**. These findings underscore the model’s robustness and ability to identify subset antibacterial compounds from large commercial libraries. By focusing on compounds with high-confidence predictions (in this case > 0.5), we can potentially prioritize candidates for further experimental validation and development or subset more specific novel antimicrobial libraries, potentially accelerating the discovery of new treatments for AMR. The predictions for all the compounds in the Enamine library can be found on Zenodo (see **Section 4.8: Data and Software Availability**).

**Figure 9:**
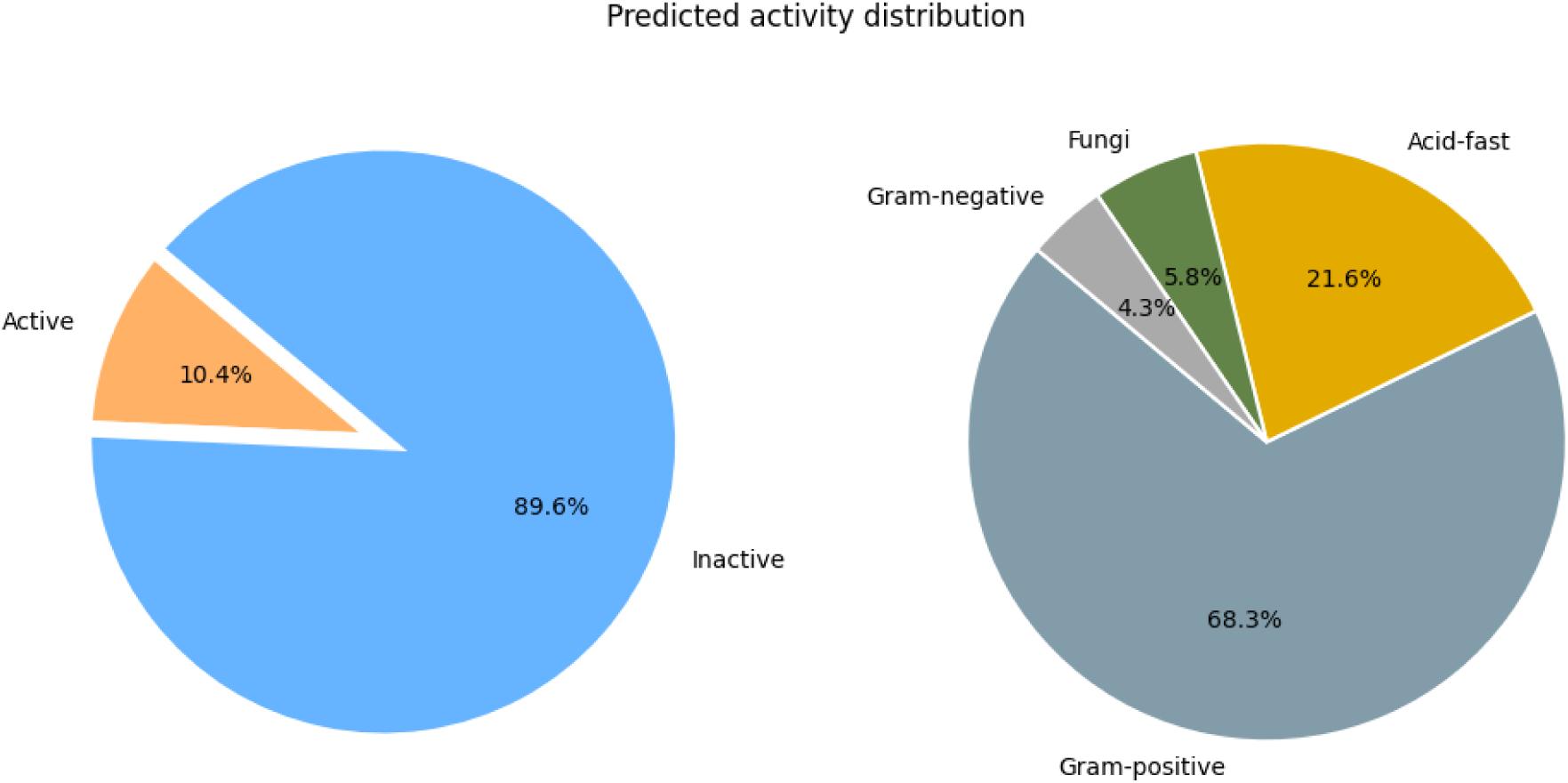
Overview of hits in Enamine Antibacterial Library based on model predictions. The left pie chart summarizes the overall predictions, while the right chart shows the distribution of predicted active compounds (10%) in the different pathogen classes.

## 3. Discussion and conclusion

Antimicrobial resistance (AMR) has been exponentially emerging as a major threat and is projected to be the leading cause of death by 2050 [46–48]. A recent study highlighted that about 5 million deaths per year are associated with AMR, with an increasing burden across low and middle-income countries [49–51]. With this growing crisis, there is an urgent need for advancements in delivering potential drugs to tackle antibiotic resistance. To assist this effort, we have developed machine learning models designed to streamline and accelerate the AMR drug discovery workflow, thereby supporting researchers in identifying effective treatments more efficiently.

Previous attempts in this direction were limited by their scope, such as focusing solely on antibacterial peptide prediction [4, 52] or bacterial species specificity [53, 54]. Additionally, such studies often demonstrate a limited use of training datasets from a single source or in-house data. Our approach aims to overcome these limitations by generating a comprehensive dataset covering a broader scope, enhancing the potential to discover effective antimicrobial treatments. We illustrated that integrating multiple public dataset collections is possible using FAIR principles. This approach ensured the consistent use of ontologies and controlled vocabularies, resulting in a medicinal chemistry-driven small-molecule bioactivity graph, the AntiMicrobial-KG. The KG consisted of experimental bioassay data for 81,490 compounds stored in a property-based graph format. The bioactivity endpoints were tested across 1,373 species, categorized into four broad pathogen classes: Gram-positive, Gram-negative, Acid-fast, and Fungi. Through the KG, we deciphered certain trends between the compounds tested for microbes and fungi. As expected, a larger number of compounds were tested against Gram-positive (∼21,000 compounds) and Gram-negative (∼9,000 compounds) pathogens compared to the Acid-fast pathogens (∼6,000 compounds). Moreover, the KG revealed a substantial presence of compounds with Michael acceptors, which have been previously known to exhibit antimicrobial and antifungal properties [40–42]. Additionally, a dominance of specific chemical classes was observed. For instance, avermectins and sesquiterpenoids were predominantly found in compounds tested against Fungi, while azo and azoxy alkaloids were prevalent in compounds tested for Gram-positive pathogens.

Following this, we showcased one potential use of the AntiMicrobial-KG in the context of the current trend of ML-based predictions. To do so, firstly, we converted the compounds in AntiMicrobial-KG into the descriptive dataset using chemoinformatics techniques. Chemoinformatics offers more than 4,000 descriptors based on 2D and 3D representations of a compound [55]. However, many of these descriptors are highly correlated, necessitating strategic selection of the most representative ones or, ideally, the most interpretable ones. Given the debate about the superiority of 2D descriptors over 3D ones, we chose to avoid 3D descriptor-based representation [56]. Complementary to compound classical descriptors, compound structural features can also be represented through fingerprints [57]. Hence, we decided to investigate multiple fingerprinting methods based on their compactness and ability to represent complementary information such as physicochemical properties, structural connectivity, and pharmacophoric contents. Our exploration of the ML-based prediction approaches yielded various conclusions. Firstly, addressing the imbalances in compounds tested across the five activity groups (Gram-positive, Gram-negative, Acid-Fast, Fungi, and Inactive) with the SMOTE technique significantly improved model predictions. Secondly, training models through systematic, progressive selection criteria assist in identifying the best model for a specific use case. Historically, ML models were trained based on community recognition (such as random forests) or black-box models (such as deep neural networks), with a few researchers comparing across a range of models. For instance, we found an average of 6-fold difference between the least-performant model (Naive Bayes) and the top-performing one (Random Forest), thus assisting us in selecting the best model from the cohort. Analogously, we used several molecular fingerprints to determine the best model-fingerprint combination. For our use case, the MHFP6-Random forest combination performed the best of all possible combinations. Lastly, while classical structural and pharmacophoric-based fingerprints outperform physicochemical properties, training models on the physicochemical representation of compounds can offer valuable insights into drug development and optimization [58]. For instance, our ChemPhys-Random Forest model highlighted hydrogen bond donor and LogP as key characteristics influencing antifungal and antibacterial activity. This finding aligns with existing research, reinforcing its relevance [59]. Additionally, we found that the number of aliphatic rings in a compound could contribute positively toward Gram-positive activity and negatively towards Gram-negative activity. Lastly, we demonstrated the applicability of our model predictions on two compound libraries commonly used in drug discovery: the EU-OPENSCREEN library and the Enamine Antibacterial collection. Our goal was to showcase the practical use of ML models for the preliminary screening of compound activities across libraries. The strategy involved improving filtering options, allowing users to select molecules with the highest probabilities of activity, thus reducing screening costs. Additionally, this approach can help commercial library vendors enhance their collections by including more active molecules. More ambitiously, we aimed to confirm and highlight key chemical features in the screened compounds that might positively distinguish active from inactive compounds for each pathogen class. Any novel insights in this area would be valuable, given that the training set used here represents one of the most extensive chemical spaces ever assembled from public sources to our knowledge. Moreover, these general predictions can be further refined to target specific bacterial species using existing ML models [60].

Despite the promising results, our current approach has, however, limitations at multiple stages, from data processing to modeling and prediction steps. First, the pathogen classes assigned to the compounds were based on the highest recorded MIC activity, meaning each compound was linked to only one pathogen class. This association implies high risks and could be misleading, as many “broad-spectrum” compounds show activity across multiple pathogen classes. Additionally, the majority of activities analyzed were driven by cellular responses, which were biased by differences in cellular permeability across pathogens, potentially skewing the results in an untracked way. Another challenge lies in data collection assembly. While some sources, such as ChEMBL, ensure data quality through manual curation, others, like SPARK and CO-ADD, may employ quality control methods of inhomogeneous levels. These distinct approaches might not necessarily be synergistic, thus yielding variations in overall data quality. On the modeling side, the lack of benchmark and validation datasets of the same size also posed limitations. The absence of structural diversity in the model’s validation sets may have hampered the model’s ability to produce more generalizable results. Nevertheless, the dynamic nature of antibiotic research, with its ever-growing number of published datasets combined with the predictive models we present here, offers a valuable tool for investigators to test and predict outcomes on their datasets. Lastly, while the inference of physicochemical “trend rules” through feature importance analysis is tied to the training dataset, one could doubt that the model’s findings reflect real inherent patterns in the data but only patterns present in the training set. However, this is similar to other widely accepted rules, such as Lipinski’s [61] or Veber’s [62], which were also derived from such data-driven inferences. In reality, being the chemical space size from which our “trend rules” are derived, the largest published dataset to date, the approach offers a more robust foundation for future research than any former attempts in this direction. Last but not least, our approach in training ML models using chemical fingerprints allows interested users, academic or commercial, to contribute to the future development of the models using the shared website for predictions of their molecules while preserving compound structural information. Moreover, with the current model, researchers could quickly test (*in-silico*) chemical libraries for antimicrobial activity and subsequently either validate the results from the model or screen new chemical space and provide the results for training the model for better accuracy in the future.

## 4. Methods

### 4.1. Aggregation of antibacterial experimental data

We created an Antimicrobial Resistant Knowledge Graph (AntiMicrobial-KG), an exhaustive data warehouse of experimentally validated antibacterial chemicals covering Gram-positive, Gram-negative, acid-fast bacteria and fungi. The construction of the AntiMicrobial-KG involved collecting minimum inhibitory concentration (MIC) data from three public data resources: CO-ADD [63], ChEMBL [64], and SPARK [65] (**Table 2**). Since each resource used unique identifiers for chemicals and the bacterial species they were tested on, we implemented a two-step process to harmonize the dataset. Initially, we manually classified pathogens into four classes: Gram-positive, Gram-negative, Acid-fast, and Fungi. Subsequently, we standardized the chemicals to extract database identifiers and their corresponding representations in the form of SMILES, InChIKeys, and InChI. We selected only those chemical-bacterial pairs that fulfilled two conditions: a) exact MIC50 value (i.e., those results with greater than or less than signs were omitted, and only those with exact values with sign (=) or less than equal to sign (≤) retained) and b) experiments reported with standard result units (i.e., µg/mL) were selected. The original source of the tested chemicals was retained, allowing the traceability of chemical-bacteria pairs to their origin. Moreover, to enable efficient comparison across these different experiments and resources, we standardized experimental values into a logarithmic scale metric similar to that implemented within ChEMBL (known as the pChEMBL value) [64]. A pChEMBL cut-off of greater than 0 was used to avoid certain erroneous activity results like negative values. This approach facilitated consistent and comparable analysis of antimicrobial resistance data across diverse datasets and experimental conditions.

**Table 2:** Number of the chemicals collected from the different data resources in the four categories (Gram-positive, Gram-negative, Acid-fast, and Fungi). It is key to note that the same chemical can be analyzed in multiple strains, and hence, the final total (i.e. # Unique total) of the chemicals is calculated based on distinct InChiKey representations. Chemicals in each of these categories may be active or inactive against the pathogen class.

| Resource Name | # Gram-positive | # Gram-negative | # Acid-fast | # Fungi | # Unique Total |
| --- | --- | --- | --- | --- | --- |
| ChEMBL (v.34) | 43,976 | 33,310 | 17,974 | 11,488 | 66,575 |
| CO-ADD | 147 | 113 | - | 112 | 286 |
| SPARK | 3,085 | 13,826 | 162 | 2 | 14,629 |

#### CO-ADD

Led by the University of Queensland, Australia, CO-ADD (http://db.co-add.org/) is a collaborative crowdsourcing approach aimed at advancing antibiotic drug discovery [63]. The repository is strategically curated with screening experimental data collected for ESKAPE pathogens (*Enterococcus faecium*, *Staphylococcus aureus*, *Klebsiella pneumoniae*, *Acinetobacter baumannii*, *Pseudomonas aeruginosa,* and *Enterobacter species*) along with various fungal strains [66]. This bacterial profiling dataset covers over 9,000 small molecules and peptide-based chemicals. CO-ADD is distinguished by its experimental approach, where the same chemical is tested across its entire strain library. This allows for in-depth chemical-strain specificity studies, a feature not commonly found in other resources where chemicals are typically tested for a single strain or strain class. As a result, the AntiMicrobial-KG integrates 286 chemicals from this resource that fit the previously described conditions (i.e., experimental endpoints with exact values and pChEMBL > 0).

#### ChEMBL

ChEMBL version 34 (https://www.ebi.ac.uk/chembl/) is a large-scale database repository for bioactivity data, focusing on small molecules and their effects on biological entities, including proteins, cell lines, and entire organisms [64]. The database provides detailed information on the resulting bioactivities of these interactions. ChEMBL is one of the most comprehensive resources for bioassays, capturing a wide range of data, including binding (B), functional (F), adsorption (A), distribution (D), metabolism (M), excretion (E), and toxicity (T) related chemical bioactivity recorded in literature and patent documents. With over 1.6 million bioassays from multiple species documented, ChEMBL serves as a critical resource for integrating extensive bioactivity data into the AntiMicrobial-KG. From this repository, we selected approximately 63,000 data points involving around 66,000 chemicals in 1,317 strains.

#### SPARK

The Shared Platform for Antibiotic Research (SPARK), now integrated and maintained by the CO-ADD community, was initially created by the Pew Charitable Trusts to expand research around antibiotics targeting Gram-negative bacteria [65]. Through this initiative, stakeholders from industry, academia, and government collaborated to develop an open platform for data sharing with the support of Collaborative Drug Discovery (CDD). Industrial partners, including Novartis and Merck, contributed published and unpublished experimental data to CDD, where domain experts, including microbiologists, curated the data to ensure consistency and establish evidence links for the recorded activities. The curation process involved harmonizing the data to maintain uniformity and reliability. The curated data was then reintegrated into CDD, creating a valuable resource for researchers and drug developers. Over time, SPARK expanded beyond Gram-negative bacterial strains to include a broader range of bacterial and fungal strains. For the SPARK data, an additional pre-processing step was performed to ensure compliance with tidy principles, wherein the standard relation and standard value are present in two columns [67]. Within the AntiMicrobial-KG, we incorporated data for approximately 14,629 chemicals from SPARK.

Pooling together the public bioassay resources, we constructed our AntiMicrobial-KG. As illustrated in **Figure 10**, some chemicals appeared in multiple databases, necessitating an additional standardization step to deduplicate chemicals tested on the same strain class. To address this, we retained only the chemical with the assay value closest to the median of all reported pMIC values. This process reduced the collected dataset from 81,486 to 74,202 chemicals. Notably, all assays in the final collection possessed non-zero values, a characteristic prominently observed in SPARK datasets. In addition to storing chemical activity data, we also incorporated chemical classifications using the NPClassifier (https://npclassifier.ucsd.edu/) [68]. This tool employs a chemical classification ontology to annotate properties, such as the pathways (for e.g., alkaloids, terpenoids, etc.) and superclasses (for e.g., glycerophospholipids, fatty amines, beta-lactams, etc.) in which the chemicals are involved. This ensured that the AntiMicrobial-KG not only captures bioactivity data but also insights into the functional roles of the chemicals, thereby enhancing overall utility.

**Figure 10:**
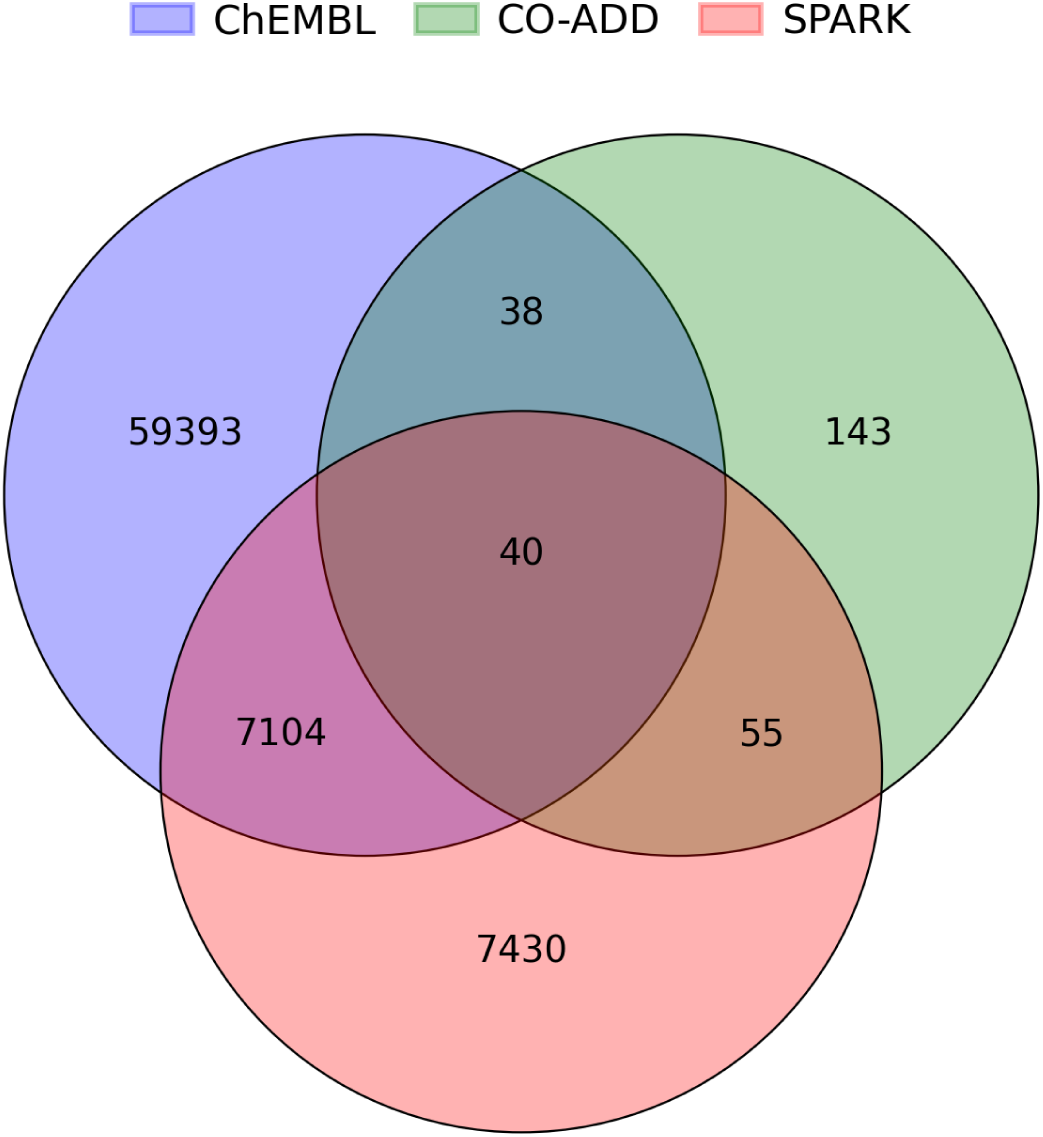
Set diagram showing the overlap between the chemicals found across the three data resources. A total of 40 chemicals are reported in all databases, while other chemicals remain unique to their original data repositories.

### 4.2. Fingerprint generation for model training

To train the models, chemical structures from the AntiMicrobial-KG are encoded into fingerprint vectors. A fingerprint is a classic representation of a chemical, serving as a molecular descriptor that relies on factors such as molecular connectivity, physicochemical properties, and functional group annotations. For this purpose, we employed the open-source tool RDKit (https://www.rdkit.org/). Utilizing RDKit, we converted 2D chemical structures into various classic fingerprints, including Morgan or Extended-connectivity fingerprints with a bond order of 4 (ECFP4) [69], topological or RDKit fingerprint with 1-7 atoms (RDKit), Molecular ACCess System (MACCS) [70], and 2D Pharmacophore fingerprint ErG [71]. Each of these fingerprints represents a one-hot encoding vector of bit sizes 1024, 2048, 167, and 315, respectively. Additionally, we incorporated the MinHash fingerprint, up to six bonds (MHF6, 2048 bits), for its proven efficacy in chemical retrieval [72]. Alongside these binary vector fingerprints that provide limited information about the chemical features, we added a physicochemical fingerprint (ChemPhys) with 29 molecular properties. The main benefit of this fingerprint was its ease of interpretability, which allowed for insights into molecular optimization strategy. These properties include generic descriptors such as SLogP, surface molecular resonance (SMR), Labute accessible surface area (ASA), polar surface area (TPSA), molecular weight (MW), number of Lipinski hydrogen bond donors (HBD) and acceptors (HBA), number of rotatable bonds, number of HBD, number of HBA, number of amide bonds, number of heteroatoms, number of heavy atoms, the total number of atoms, and various counts of rings (total, aromatic, saturated and aliphatic). Specific descriptors included fragment complexity, rotatable bond count (terminal and non-terminal), number of bridgehead carbon atoms shared between rings, number of stereocenters (specified and unspecified), and counts of heterocycles (aromatic and saturated) and carbocycles (aromatic, saturated, and aliphatic).

To ensure the robustness of all feature columns (i.e., the bit vector representing the chemical fingerprint), we conducted a check for null variance in each feature across the chemicals. The reason for performing a null variance check is to reduce noise for model training and to confirm that every feature provided meaningful information, thus contributing to the chemical characterization for the model’s understanding. By integrating these diverse fingerprinting techniques, we created a comprehensive and detailed representation of the chemical structures, facilitating effective model training and subsequent predictions.

### 4.3. Train and test datasets

For training and evaluating our machine learning (ML) model, we split the chemical data in AntiMicrobial-KG into train and test sets. Since our goal was to predict the activity of chemicals, the dataset was organized into chemical-pathogen class pairs. Compounds were first classified as active (pChEMBL > 5) or inactive (pChEMBL ≤ 5) based on their pChEMBL values. For those in the active category, pathogen class selectivity was determined using a “best-of-four” approach, where the pathogen class (Gram-positive, Gram-negative, fungi, or acid-fast) with the highest potency, as indicated by its pChEMBL value, was selected for each chemical. If the compound was found to be inactive against all pathogens, it was labeled in the “inactive” category.

The initial distribution of chemicals across the five categories revealed class imbalances: 21,148 chemicals in Gram-positive, 9,083 in Gram-negative, 7,631 in fungi, 5,845 in acid-fast and 15,654 in inactive classes. To address this imbalance and enhance the robustness of our ML model, we employed an oversampling strategy during dataset splitting. Specifically, we used the Synthetic Minority Over-sampling Technique (SMOTE), which involves either undersampling of the majority class (in this case, the Gram-positive class) or oversampling of the minority class (in this case, the fungi class) to create synthetic sampled to balance the class [73]. This resulted in a balanced dataset with 21,148 chemical-pathogen class pairs in each class.

For our training-testing splits, we allocated approximately 80% of chemical-pathogen pairs to the training set and the remaining 20% to the test set. Importantly, SMOTE was applied only to the training data, ensuring that the test set remained an accurate representation of the real-world data distribution. This approach allowed us to train more robust and reliable ML models capable of predicting chemical activities across different pathogen classes.

### 4.4. Development and evaluation of machine learning models

For this multiclass classification task, we developed a reproducible pipeline to test multiple models and optimize the best-performing one. This pipeline was designed with versatility in mind, allowing it to be applied to any use case, provided the data is available in the requisite format. We began by leveraging PyCaret’s (https://pycaret.org/) model comparison pipeline to streamline the selection process for the best-performing ML models from a cohort. In this pipeline, we trained six classic ML models: Naive Bayes (NB), Logistic regression (LR), Light Gradient Boosting Machine (LightGBM), Decision Tree (DT), Random Forest (RF), and eXtreme Gradient Boosting (XGBoost) with a 5-fold cross-validation strategy. The choice of these models was driven by their transparent nature, as opposed to the black-box nature of certain advanced ML models like neural networks. For training these models, the initial training data (80%) was further split into train (60%) and validation or hold-out set (20%). The trained models were ranked based on their performance in predicting the hold-out set, using the Cohen-Kappa score as the performance metric. It is important to note that PyCaret’s model training pipeline internally applies various data preprocessing steps, including imputation, scaling, and normalization. To further refine the best model, we performed an additional optimization strategy using hyperparameter optimization (HPO) with Optuna (https://optuna.org/). This HPO pipeline involved 15 trial runs aiming to maximize the Cohen-Kappa score. Similar to PyCaret, the score for optimization was determined from the 5-fold cross-validation strategy. This two-layered approach (model selection and model refinement) ensures the development of a robust and globally optimized model for evaluation in our multiclass classification task.

In addition to the Cohen Kappa score, we calculated other prediction metrics such as accuracy and Area Under the Curve of Receiver Operating Characteristic (AUC-ROC). These metrics were used not only to select the best model from our collection but also to compare our model with existing ones in the field of AMR. Once the optimal model was identified, it was used to predict the outcomes of the test set (the remaining 20% of the dataset), demonstrating its effectiveness and applicability. Furthermore, we used Shapley values to identify the most influential features affecting the strain class within the dataset. Identifying the top ten most influential features enhances the transparency and interpretability of the model, regardless of the algorithm employed. This robust and transparent approach ensures that our model is not only highly accurate but also interpretable, providing valuable insights into the chemical features driving antimicrobial resistance.

### 4.5. External compound libraries used for predictions and validation

Additionally, external datasets were collected to evaluate and predict the antibacterial, antifungal, and antituberculosis activity of compounds. This involved the compounds from two sources: a) the subset of Enamine compound library (with 32,000 compounds) specially optimized for AMR, the Antibacterial Library (https://enamine.net/compound-libraries/targeted-libraries/antibacterial-library) and b) EU Openscreen (EU-OS) European Chemical Biology Library (https://ecbd.eu/) with 101,024 compounds. The Enamine library was designed with privileged scaffolds from known antibacterial drugs and their underlying physicochemical properties in mind. The EU-OS library was designed based on four different chemoinformatic-rich content approaches and is not directed towards antibacterial but rather is a collection that includes a range of novel and diverse scaffolds. To distinguish these two, we call them the ECBL set (96,092 compounds) and the Bioactive set (4,927) and the results of the reports for these sets independently.

To compare the model predictions with the experimental predictions, we make use of the hit rate. The HitRate for a specific pathogen class is described as:

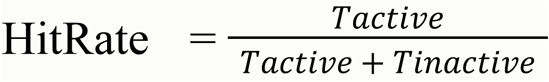

where T*_active_* represents the bioactive compounds, and T*_inactive_* denotes the inactive compounds for the specific pathogen class. For experimental results, a compound is considered active if it shows 50% or greater inhibition. This metric allows for a quantitative comparison of the hits identified by the model with the experimental “gold standard” results.

### 4.6. Experimental testing of EU-OS EBCL library

In addition to the model predictions, the library was tested in a high-content screening viability assay against seven microbial pathogens: three fungi (*C. auris* DSM21092, *C. albicans* ATCC 64124, *A. fumigatus* ATCC 46645), two Gram-positive (*S. aureus* ATCC 29213 and *E. faecalis* ATCC 29212), and two Gram-negative (*P. aeruginosa* and *E. coli*). A compound concentration of 50µM was used, and the results were reported as percentage inhibition. All results, along with the experimental protocols, can be found in the ECBD database (https://ecbd.eu/) with the following assay identifiers:

- *C. auris* DSM21092 - EOS300072
- *C. albicans* ATCC 64124 - EOS300076
- A. fumigatus ATCC 46645 - EOS300074
- *S. aureus* ATCC 29213 - EOS300078
- *E. faecalis* ATCC 29212 - EOS300080
- *P. aeruginosa* group - EOS300155
- *E. coli* ATCC 25922 - EOS300158

### 4.7. Implementation

The dataset was generated using Python version 3.9. For the generation of the chemical fingerprints and molecular scaffolds, we took advantage of RDKit as implemented in Python. The stratification of the data into train-test splits was performed using the Scikit-learn library (https://scikit-learn.org), and SMOTE was implemented using the Imbalance-learn library (https://imbalanced-learn.org). To ensure the reproducibility of the sampling procedures, the data splitting was seed-coded. The models were built in PyCaret (v.3.2.0) and Scikit-learn (v.1.3.2). Feature importance was determined using the TreeInterpretor (https://pypi.org/project/treeinterpreter/) python package. For the AntiMicrobial-KG, we provide the opportunity to generate the graph with Neo4J (https://neo4j.com/), a commercially available graph database analysis and visualization tool. The website for displaying the database information was built using Streamlit (https://streamlit.io/) and is hosted on SciLifeLab’s SERVE (https://serve.scilifelab.se/) instance.

### 4.8. Data and Software Availability

All the Python scripts used in this manuscript for model training, exploratory analysis, and KG generation are available on GitHub at https://github.com/IMI-COMBINE/broad_spectrum_prediction. The data collected with the AntiMicrobial-KG and the models can also be found on Zenodo at https://zenodo.org/records/13868088. Also, we have the AntiMicrobial-KG website at https://antimicrobial-kg.serve.scilifelab.se/ to allow users to search the database and use our pre-trained models for their compound activity prediction.

## Supporting Information

**Figure S1:** Non-promiscuous scaffolds found in AntiMicrobial-KG.

**Figure S2:** NP-likeness distribution of chemicals found in AntiMicrobial-KG.

**Figure S3:** Comparison of SMOTE vs non-SMOTE for the top two performing models.

**Figure S4:** Feature importance of MHFP6 fingerprints.

**Figure S5:** Feature importance of MHFP6 fingerprints.

**Table S1:** Summary of Ro5 violations and structural alerts in AMR-KG.

**Table S2:** Evaluation metrics on hold-out set for all chemical fingerprints across six ML models.

**Table S3:** Hyperparameters used for optimizing the top two models: Random forest and XGBoost.

**Table S4:** XGBoost model performance on the test data.

**Table S5:** Random Forest model performance on the test data.

**Table S6:** EU-OS library prediction for each pathogen class.

**Table S7:** High confidence prediction of Enamine library.

### Acknowledgments

We would like to thank the authors of the resources used in our work for making their datasets available to the scientific community. We would like to acknowledge Dr. Frederik Deroose, Dr. Anders Karlén, Dr. Axel Pahl, Dr. Maria Kuzikov and all other members of the AMR Accelerator Scientific Interest Group on Machine Learning group for their feedback for improving the manuscript. We would also like to thank the anonymous reviewers for their comments and suggestions for improving the readability of the manuscript.

## Author contribution

Y.G: Conceptualization, Methodology, Writing - Original draft preparation and Visualization; A.Z: Conceptualization, Supervision, Methodology, Writing - Original draft preparation and Visualization; L.B: Visualization; O.G: Investigation; U.B: Investigation; M.B: Investigation; M.K: Writing - Reviewing and Editing; P.G: Writing - Reviewing and Editing.

## References

1. Palmer JD, Foster KR. The evolution of spectrum in antibiotics and bacteriocins. Proceedings of the National Academy of Sciences. 2022 Sep 20;119(38):e2205407119. 10.1073/pnas.2205407119

2. Peterson E, Kaur P. Antibiotic resistance mechanisms in bacteria: relationships between resistance determinants of antibiotic producers, environmental bacteria, and clinical pathogens. Frontiers in microbiology. 2018 Nov 30;9:2928. 10.3389/fmicb.2018.02928

3. O’Neill J. Tackling drug-resistant infections globally: final report and recommendations. https://apo.org.au/node/63983

4. Murray CJ, Ikuta KS, Sharara F, Swetschinski L, Aguilar GR, Gray A, Han C, Bisignano C, Rao P, Wool E, Johnson SC. Global burden of bacterial antimicrobial resistance in 2019: a systematic analysis. The lancet. 2022 Feb 12;399(10325):629–55. 10.1016/S0140-6736(21)02724-0

5. Mestrovic T, Aguilar GR, Swetschinski LR, Ikuta KS, Gray AP, Weaver ND, Han C, Wool EE, Hayoon AG, Hay SI, Dolecek C. The burden of bacterial antimicrobial resistance in the WHO European region in 2019: a cross-country systematic analysis. The Lancet Public Health. 2022 Nov 1;7(11):e897–913. 10.1016/S2468-2667(22)00225-0

6. Essack SY, Lenglet A. Bacterial antimicrobial resistance burden in Africa: accuracy, action, and alternatives. The Lancet Global Health. 2024 Feb 1;12(2):e171–2. 10.1016/S2214-109X(23)00587-9

7. Ventola CL. The antibiotic resistance crisis: part 1: causes and threats. Pharmacy and therapeutics. 2015 Apr;40(4):277.

8. Zankari E, Hasman H, Kaas RS, Seyfarth AM, Agersø Y, Lund O, Larsen MV, Aarestrup FM. Genotyping using whole-genome sequencing is a realistic alternative to surveillance based on phenotypic antimicrobial susceptibility testing. Journal of antimicrobial chemotherapy. 2013 Apr 1;68(4):771–7. 10.1093/jac/dks496

9. Genestet C, Hodille E, Berland JL, Ginevra C, Bryant JE, Ader F, Lina G, Dumitrescu O, Lyon TB Study Group. Whole-genome sequencing in drug susceptibility testing of Mycobacterium tuberculosis in routine practice in Lyon, France. International journal of antimicrobial agents. 2020 Apr 1;55(4):105912. 10.1016/j.ijantimicag.2020.105912

10. Ellington MJ, Ekelund O, Aarestrup FM, Canton R, Doumith M, Giske C, Grundman H, Hasman H, Holden MT, Hopkins KL, Iredell J. The role of whole genome sequencing in antimicrobial susceptibility testing of bacteria: report from the EUCAST Subcommittee. Clinical microbiology and infection. 2017 Jan 1;23(1):2–2. 10.1016/j.cmi.2016.11.012

11. Su M, Satola SW, Read TD. Genome-based prediction of bacterial antibiotic resistance. Journal of clinical microbiology. 2019 Mar;57(3):10–128. 10.1128/jcm.01405-18

12. Lundstrom TS, Sobel JD. Antibiotics for gram-positive bacterial infections: vancomycin, teicoplanin, quinupristin/dalfopristin, and linezolid. Infectious Disease Clinics. 2000 Jun 1;14(2):463–74. 10.1016/S0891-5520(05)70258-0

13. Abu-Farha R, Gharaibeh L, Alzoubi KH, Nazal R, Zawiah M, Binsaleh AY, Shilbayeh SA. Awareness, perspectives and practices of antibiotics deprescribing among physicians in Jordan: a cross-sectional study. Journal of Pharmaceutical Policy and Practice. 2024 Dec 31;17(1):2378484. 10.1080/20523211.2024.2378484

14. Bdair IA, Bdair OA, Maribbay GM, Elzehiri D, Hassan ES, Tolentino AD, Emara MM, Ali EK, Abdelmeged RM, Fadul MO. Public awareness towards antibiotics use, misuse and resistance in Saudi community: A cross-sectional population survey. Journal of Applied Pharmaceutical Science. 2024 Jul 10. 10.7324/JAPS.2024.167162

15. Rodríguez-Gascón A, Solinís MÁ, Isla A. The role of PK/PD analysis in the development and evaluation of antimicrobials. Pharmaceutics. 2021 Jun 3;13(6):833. 10.3390/pharmaceutics13060833

16. Sy SK, Derendorf H. Pharmacokinetics I: PK-PD approach, the case of antibiotic drug development. Clinical pharmacology: current topics and case studies. 2016:185–217. 10.1007/978-3-319-27347-1_13

17. Sou T, Hansen J, Liepinsh E, Backlund M, Ercan O, Grinberga S, Cao S, Giachou P, Petersson A, Tomczak M, Urbas M. Model-informed drug development for antimicrobials: translational PK and PK/PD modeling to predict an efficacious human dose for apramycin. Clinical Pharmacology & Therapeutics. 2021 Apr;109(4):1063–73. 10.1002/cpt.2104

18. Lin TT, Yang LY, Lin CY, Wang CT, Lai CW, Ko CF, Shih YH, Chen SH. Intelligent de novo design of novel antimicrobial peptides against antibiotic-resistant bacteria strains. International journal of molecular sciences. 2023 Apr 5;24(7):6788. 10.3390/ijms24076788

19. Jukič M, Bren U. Machine learning in antibacterial drug design. Frontiers in Pharmacology. 2022 May 3;13:864412. 10.3389/fphar.2022.864412

20. Melo MC, Maasch JR, de la Fuente-Nunez C. Accelerating antibiotic discovery through artificial intelligence. Communications biology. 2021 Sep 9;4(1):1050. 10.1038/s42003-021-02586-0

21. Anahtar MN, Yang JH, Kanjilal S. Applications of machine learning to the problem of antimicrobial resistance: an emerging model for translational research. Journal of clinical microbiology. 2021 Jun 18;59(7):10–128. 10.1128/jcm.01260-20

22. Sakagianni A, Koufopoulou C, Feretzakis G, Kalles D, Verykios VS, Myrianthefs P. Using machine learning to predict antimicrobial resistance―a literature review. Antibiotics. 2023 Feb 24;12(3):452. 10.3390/antibiotics12030452

23. Tyers M, Wright GD. Drug combinations: a strategy to extend the life of antibiotics in the 21st century. Nature Reviews Microbiology. 2019 Mar;17(3):141–55. 10.1038/s41579-018-0141-x

24. Lv J, Liu G, Ju Y, Sun Y, Guo W. Prediction of synergistic antibiotic combinations by graph learning. Frontiers in Pharmacology. 2022 Mar 8;13:849006. 10.3389/fphar.2022.849006

25. Yang Y, Ai C, Ji Z, Chen W, Song Y, Zeng J, Duan M, Qi W, Zhang S, An Z, Lin Y. Antibiotic combinations prediction based on machine learning to multicentre clinical data and drug interaction correlation. International Journal of Antimicrobial Agents. 2024 May 1;63(5):107122. 10.1016/j.ijantimicag.2024.107122

26. Cantrell JM, Chung CH, Chandrasekaran S. Machine learning to design antimicrobial combination therapies: Promises and pitfalls. Drug Discovery Today. 2022 Jun 1;27(6):1639–51. 10.1016/j.drudis.2022.04.006

27. Chio H, Guest EE, Hobman JL, Dottorini T, Hirst JD, Stekel DJ. Predicting bioactivity of antibiotic metabolites by molecular docking and dynamics. Journal of Molecular Graphics and Modelling. 2023 Sep 1;123:108508. 10.1016/j.jmgm.2023.108508

28. Alves MJ, Froufe HJ, Costa AF, Santos AF, Oliveira LG, Osório SR, Abreu RM, Pintado M, Ferreira IC. Docking studies in target proteins involved in antibacterial action mechanisms: Extending the knowledge on standard antibiotics to antimicrobial mushroom compounds. Molecules. 2014 Jan 29;19(2):1672–84. 10.3390/molecules19021672

29. Karnati P, Gonuguntala R, Barbadikar KM, Mishra D, Jha G, Prakasham V, Chilumula P, Shaik H, Pesari M, Sundaram RM, Chinnaswami K. Performance of Novel Antimicrobial Protein Bg_9562 and In Silico Predictions on Its Properties with Reference to Its Antimicrobial Efficiency against Rhizoctonia solani. Antibiotics. 2022 Mar 8;11(3):363. 10.3390/antibiotics11030363

30. Wong F, Krishnan A, Zheng EJ, Stärk H, Manson AL, Earl AM, Jaakkola T, Collins JJ. Benchmarking AlphaFold-enabled molecular docking predictions for antibiotic discovery. Molecular systems biology. 2022 Sep;18(9):e11081. 10.15252/msb.202211081

31. Zhao F, Qiu J, Xiang D, Jiao P, Cao Y, Xu Q, Qiao D, Xu H, Cao Y. deepAMPNet: a novel antimicrobial peptide predictor employing AlphaFold2 predicted structures and a bi-directional long short-term memory protein language model. PeerJ. 2024 Jul 19;12:e17729. 10.7717/peerj.17729

32. Feretzakis G, Sakagianni A, Loupelis E, Kalles D, Skarmoutsou N, Martsoukou M, Christopoulos C, Lada M, Petropoulou S, Velentza A, Michelidou S. Machine learning for antibiotic resistance prediction: A prototype using off-the-shelf techniques and entry-level data to guide empiric antimicrobial therapy. Healthcare informatics research. 2021 Jul 31;27(3):214–21. 10.4258/hir.2021.27.3.214

33. Weis CV, Jutzeler CR, Borgwardt K. Machine learning for microbial identification and antimicrobial susceptibility testing on MALDI-TOF mass spectra: a systematic review. Clinical Microbiology and Infection. 2020 Oct 1;26(10):1310–7. 10.1016/j.cmi.2020.03.014

34. Carracedo-Reboredo P, Liñares-Blanco J, Rodríguez-Fernández N, Cedrón F, Novoa FJ, Carballal A, Maojo V, Pazos A, Fernandez-Lozano C. A review on machine learning approaches and trends in drug discovery. Computational and structural biotechnology journal. 2021 Jan 1;19:4538–58. 10.1016/j.csbj.2021.08.011

35. Ivanenkov YA, Zhavoronkov A, Yamidanov RS, Osterman IA. Identification of novel antibacterials using machine learning techniques, Front. Pharmacol. 2019;10(913):10–3389. 10.3389/fphar.2019.00913

36. Fjell CD, Jenssen H, Hilpert K, Cheung WA, Panté N, Hancock RE, Cherkasov A. Identification of novel antibacterial peptides by chemoinformatics and machine learning. Journal of medicinal chemistry. 2009 Apr 9;52(7):2006–15. 10.1021/jm8015365

37. Nguyen M, Brettin T, Long SW, Musser JM, Olsen RJ, Olson R, Shukla M, Stevens RL, Xia F, Yoo H, Davis JJ. Developing an in silico minimum inhibitory concentration panel test for Klebsiella pneumoniae. Scientific reports. 2018 Jan 11;8(1):421. 10.1038/s41598-017-18972-w

38. Gurvic D, Leach AG, Zachariae U. Data-driven derivation of molecular substructures that enhance drug activity in gram-negative bacteria. Journal of Medicinal Chemistry. 2022 Apr 15;65(8):6088–99. 10.1021/acs.jmedchem.1c01984

39. Liang ST, Chen C, Chen RX, Li R, Chen WL, Jiang GH, Du LL. Michael acceptor molecules in natural products and their mechanism of action. Frontiers in Pharmacology. 2022 Nov 2;13:1033003. 10.3389/fphar.2022.1033003

40. Sherzad Othman S. Synthesis of Novel Michael Adducts and Study of their Antioxidant and Antimicrobial Activities. Chemical Review and Letters. 2022 Aug 1;5(4):226–33. 10.22034/CRL.2022.350315.1176

41. Strharsky T, Pindjakova D, Kos J, Vrablova L, Smak P, Michnova H, Gonec T, Hosek J, Oravec M, Jendrzejewska I, Cizek A. Trifluoromethylcinnamanilide michael acceptors for treatment of resistant bacterial infections. International Journal of Molecular Sciences. 2022 Dec 1;23(23):15090. 10.3390/ijms232315090

42. Lee KM, Le P, Sieber SA, Hacker SM. Degrasyn exhibits antibiotic activity against multi-resistant Staphylococcus aureus by modifying several essential cysteines. Chemical communications. 2020;56(19):2929–32. 10.1039/C9CC09204H

43. Ciura K, Fedorowicz J, Andrić F, Žuvela P, Greber KE, Baranowski P, Kawczak P, Nowakowska J, Bączek T, Sączewski J. Lipophilicity determination of antifungal isoxazolo [3, 4-b] pyridin-3 (1 H)-ones and their N1-substituted derivatives with chromatographic and computational methods. Molecules. 2019 Nov 26;24(23):4311. 10.3390/molecules24234311

44. Kokot, M., Weiss, M., Zdovc, I., Senerovic, L., Radakovic, N., Anderluh, M., Minovski, N. and Hrast, M., 2023. Amide containing NBTI antibacterials with reduced hERG inhibition, retained antimicrobial activity against gram-positive bacteria and in vivo efficacy. European journal of medicinal chemistry, 250, p.115160. 10.1016/j.ejmech.2023.115160

45. Limwongyut J, Moreland AS, Nie C, Read de Alaniz J, Bazan GC. Amide Moieties Modulate the Antimicrobial Activities of Conjugated Oligoelectrolytes against Gram-negative Bacteria. ChemistryOpen. 2022 Feb;11(2):e202100260. 10.1002/open.202100260

46. Tang KW, Millar BC, Moore JE. Antimicrobial resistance (AMR). British Journal of Biomedical Science. 2023;80:11387. 10.3389/bjbs.2023.11387

47. Lancet T. Antimicrobial resistance: an agenda for all. Lancet (London, England). 2024 May 23:S0140–6736. 10.1016/S0140-6736(24)01076-6

48. Fongang H, Mbaveng AT, Kuete V. Global burden of bacterial infections and drug resistance. *In* Advances in Botanical Research 2023 Jan 1 (Vol. 106, pp. 1–20). Academic Press. 10.1016/bs.abr.2022.08.001

49. Brüssow H. The antibiotic resistance crisis and the development of new antibiotics. Microbial Biotechnology. 2024 Jul;17(7):e14510. 10.1111/1751-7915.14510

50. Bournez C, Riool M, de Boer L, Cordfunke RA, de Best L, van Leeuwen R, Drijfhout JW, Zaat SA, van Westen GJ. CalcAMP: a new machine learning model for the accurate prediction of antimicrobial activity of peptides. Antibiotics. 2023 Apr 7;12(4):725. 10.3390/antibiotics12040725

51. Bajiya N, Choudhury S, Dhall A, Raghava GP. AntiBP3: A Method for Predicting Antibacterial Peptides against Gram-Positive/Negative/Variable Bacteria. Antibiotics. 2024 Feb 8;13(2):168. 10.3390/antibiotics13020168

52. Li JT, Wei YW, Wang MY, Yan CX, Ren X, Fu XJ. Antibacterial activity prediction model of traditional Chinese medicine based on combined data-driven approach and machine learning algorithm: constructed and validated. Frontiers in Microbiology. 2021 Nov 22;12:763498. 10.3389/fmicb.2021.763498

53. Nsubuga M, Galiwango R, Jjingo D, Mboowa G. Generalizability of machine learning in predicting antimicrobial resistance in E. coli: a multi-country case study in Africa. BMC genomics. 2024 Mar 18;25(1):287. 10.1186/s12864-024-10214-4

54. Todeschini R, Consonni V. Molecular descriptors for chemoinformatics. John Wiley & Sons; 2009 Oct 30. ISBN:9783527318520

55. Bahia MS, Kaspi O, Touitou M, Binayev I, Dhail S, Spiegel J, Khazanov N, Yosipof A, Senderowitz H. A comparison between 2D and 3D descriptors in QSAR modeling based on bio-active conformations. Molecular Informatics. 2023 Apr;42(4):2200186. 10.1002/minf.202200186

56. Duan J, Dixon SL, Lowrie JF, Sherman W. Analysis and comparison of 2D fingerprints: insights into database screening performance using eight fingerprint methods. Journal of Molecular Graphics and Modelling. 2010 Sep 1;29(2):157–70. 10.1016/j.jmgm.2010.05.008

57. Richter MF, Hergenrother PJ. The challenge of converting Gram-positive-only compounds into broad-spectrum antibiotics. Annals of the New York Academy of Sciences. 2019 Jan;1435(1):18–38. 10.1111/nyas.13598

58. Yuan G, Guan Y, Yi H, Lai S, Sun Y, Cao S. Antibacterial activity and mechanism of plant flavonoids to gram-positive bacteria predicted from their lipophilicities. Scientific reports. 2021 May 18;11(1):10471. 10.1038/s41598-021-90035-7

59. Boulaamane Y, Molina Panadero I, Hmadcha A, Atalaya Rey C, Baammi S, El Allali A, Maurady A, Smani Y. Antibiotic discovery with artificial intelligence for the treatment of Acinetobacter baumannii infections. Msystems. 2024 May 3:e00325–24. 10.1128/msystems.00325-24

60. Badura A, Krysiński J, Nowaczyk A, Buciński A. Application of artificial neural networks to prediction of new substances with antimicrobial activity against Escherichia coli. Journal of Applied Microbiology. 2021 Jan 1;130(1):40–9. 10.1111/jam.14763

61. Lipinski CA. Lead-and drug-like compounds: the rule-of-five revolution. Drug discovery today: Technologies. 2004 Dec 1;1(4):337–41. 10.1016/j.ddtec.2004.11.007

62. Veber DF, Johnson SR, Cheng HY, Smith BR, Ward KW, Kopple KD. Molecular properties that influence the oral bioavailability of drug candidates. Journal of medicinal chemistry. 2002 Jun 6;45(12):2615–23. 10.1021/jm020017n

63. Cooper MA. A community-based approach to new antibiotic discovery. Nature Reviews Drug Discovery. 2015 Sep;14(9):587–8. 10.1038/nrd4706

64. Gaulton A, Bellis LJ, Bento AP, Chambers J, Davies M, Hersey A, Light Y, McGlinchey S, Michalovich D, Al-Lazikani B, Overington JP. ChEMBL: a large-scale bioactivity database for drug discovery. Nucleic acids research. 2012 Jan 1;40(D1):D1100–7. 10.1093/nar/gkr777

65. Thomas J, Navre M, Rubio A, Coukell A. Shared platform for antibiotic research and knowledge: A collaborative tool to SPARK antibiotic discovery. ACS infectious diseases. 2018 Sep 21;4(11):1536–9. 10.1021/acsinfecdis.8b00193

66. Semenov VV, Raihstat MM, Konyushkin LD, Semenov RV, Blaskovich MA, Zuegg J, Elliott AG, Hansford KA, Cooper MA. Antimicrobial screening of a historical collection of over 140 000 small molecules. Mendeleev Communications. 2021 Jul 1;31(4):484–7. 10.1016/j.mencom.2021.07.015

67. Wickham H. Tidy data. Journal of statistical software. 2014 Sep 12;59:1–23. 10.18637/jss.v059.i10

68. Kim HW, Wang M, Leber CA, Nothias LF, Reher R, Kang KB, Van Der Hooft JJ, Dorrestein PC, Gerwick WH, Cottrell GW. NPClassifier: a deep neural network-based structural classification tool for natural products. Journal of Natural Products. 2021 Oct 18;84(11):2795–807. 10.1021/acs.jnatprod.1c00399

69. Rogers D, Hahn M. Extended-connectivity fingerprints. Journal of chemical information and modeling. 2010 May 24;50(5):742–54. 10.1021/ci100050t

70. Durant JL, Leland BA, Henry DR, Nourse JG. Reoptimization of MDL keys for use in drug discovery. Journal of chemical information and computer sciences. 2002 Nov 25;42(6):1273–80. 10.1021/ci010132r

71. Stiefl N, Watson IA, Baumann K, Zaliani A. ErG: 2D pharmacophore descriptions for scaffold hopping. Journal of chemical information and modeling. 2006 Jan 23;46(1):208–20. 10.1021/ci050457y

72. Probst D, Reymond JL. A probabilistic molecular fingerprint for big data settings. Journal of cheminformatics. 2018 Dec;10:1–2. 10.1186/s13321-018-0321-8

73. Chawla NV, Bowyer KW, Hall LO, Kegelmeyer WP. SMOTE: synthetic minority over-sampling technique. Journal of artificial intelligence research. 2002 Jun 1;16:321–57. 10.1613/jair.953

