## Supporting Information for "Predicting antimicrobial class specificity of small molecules using machine learning"

**Figures**

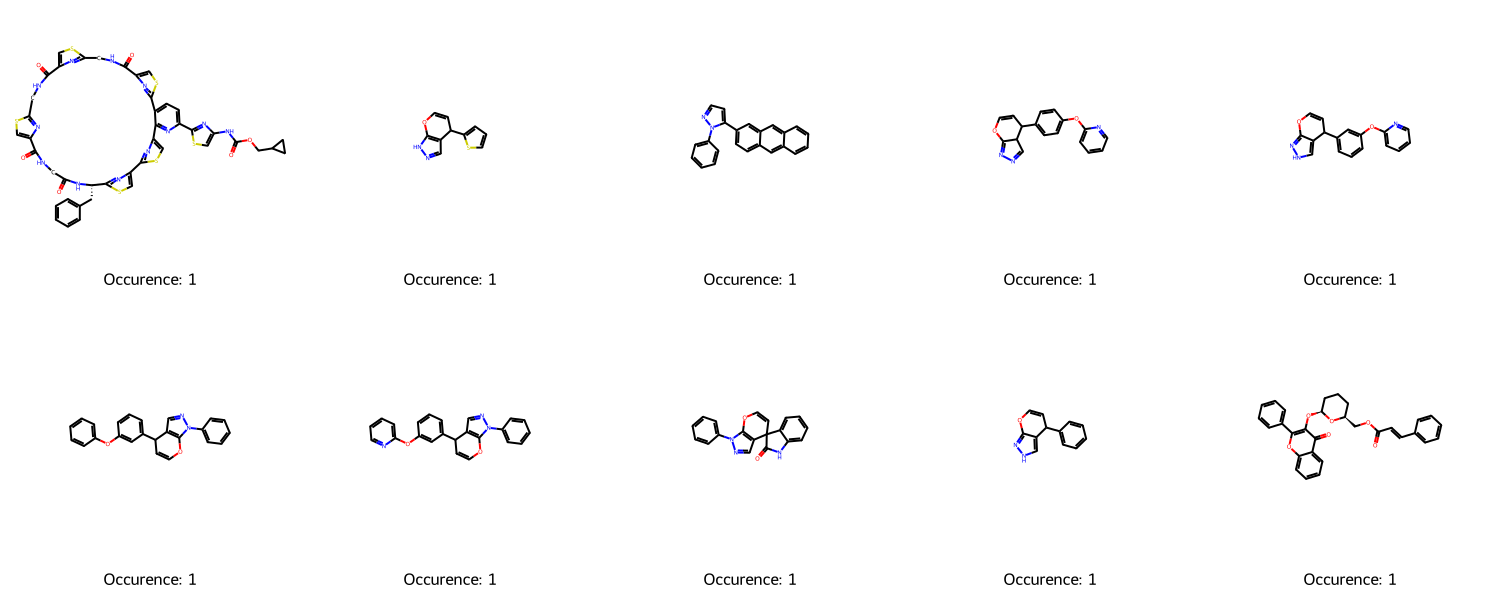

**Figure S1: Non-promiscuous scaffolds found in AntiMicrobial-KG.**

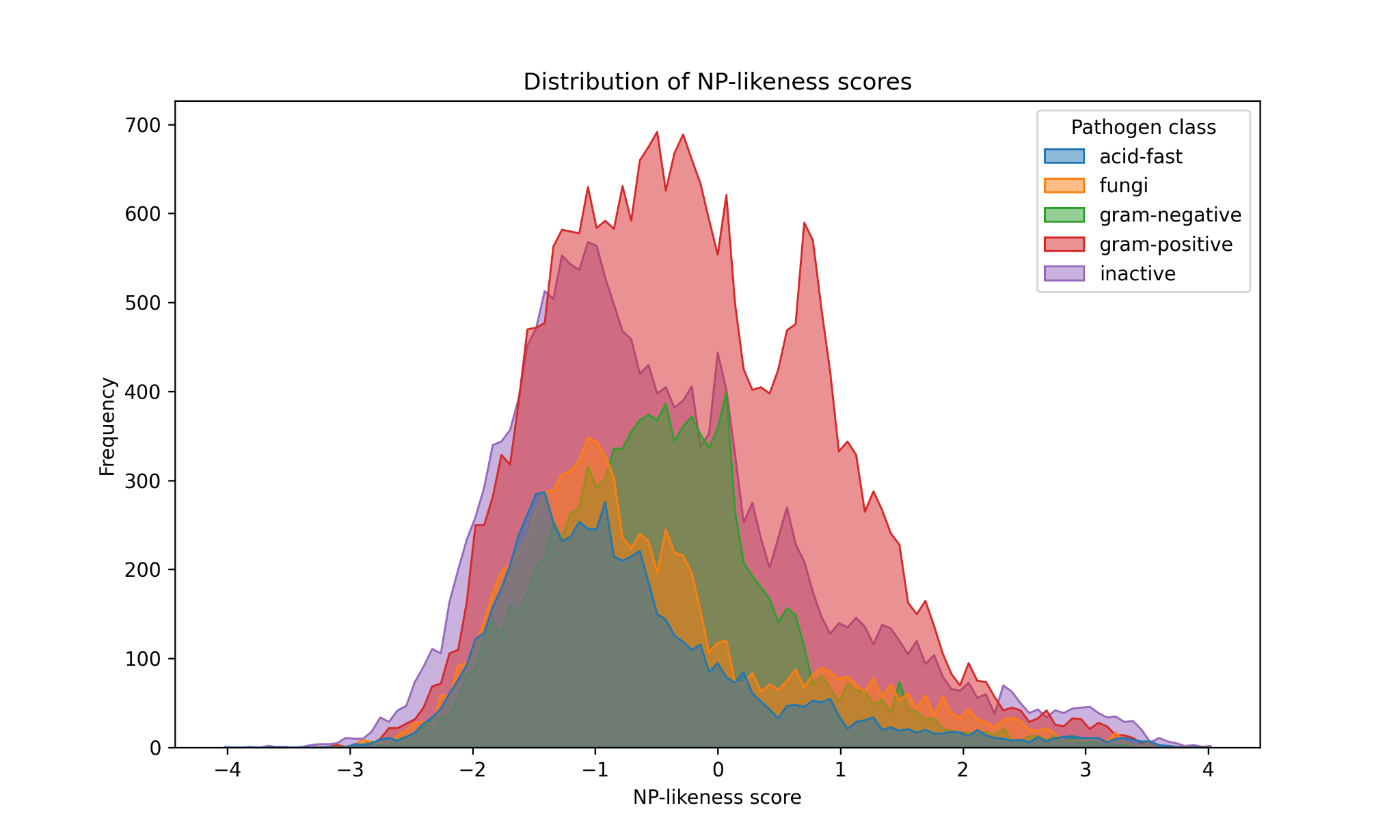

**Figure S2: NP-likeness distribution of chemicals found in AntiMicrobial-KG.** Chemicals are considered to be synthetic with an NP-likeness score < -1 and natural products with an NP-likeness score > 1.

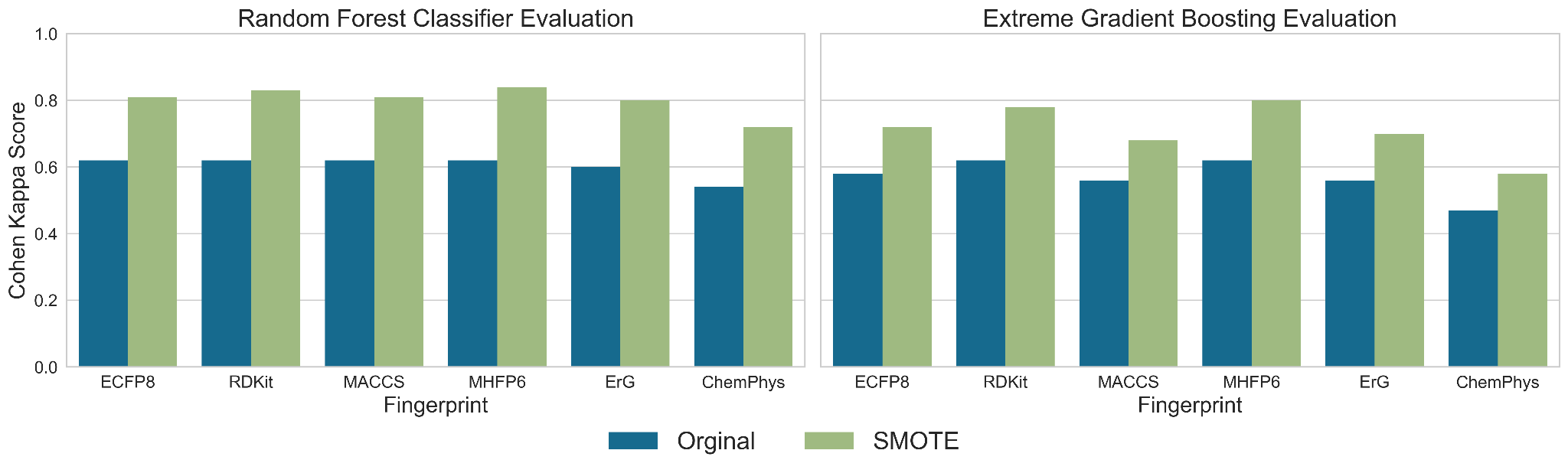

**Figure S3: Comparison of SMOTE vs non-SMOTE for the top two performing models (RF and XGBoost).** For each fingerprint, the SMOTE-based model training outperforms the classic model training.

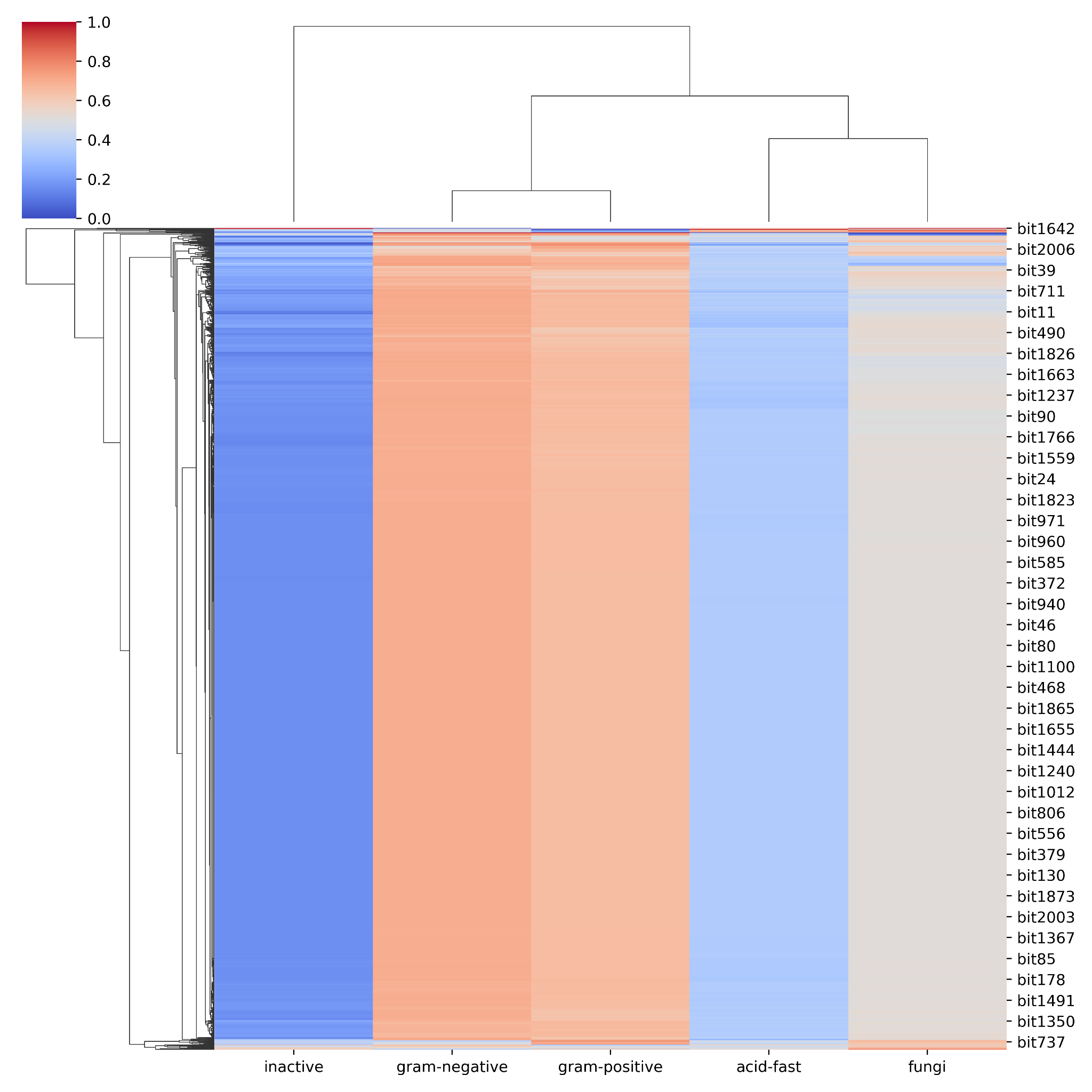

**Figure S4: Feature importance of MHFP6 fingerprints.** The rows represent the MHFP6 bits, and the columns represent the five label classes for the model. The colours in the heatmap denote the importance of the feature in the respective label class.

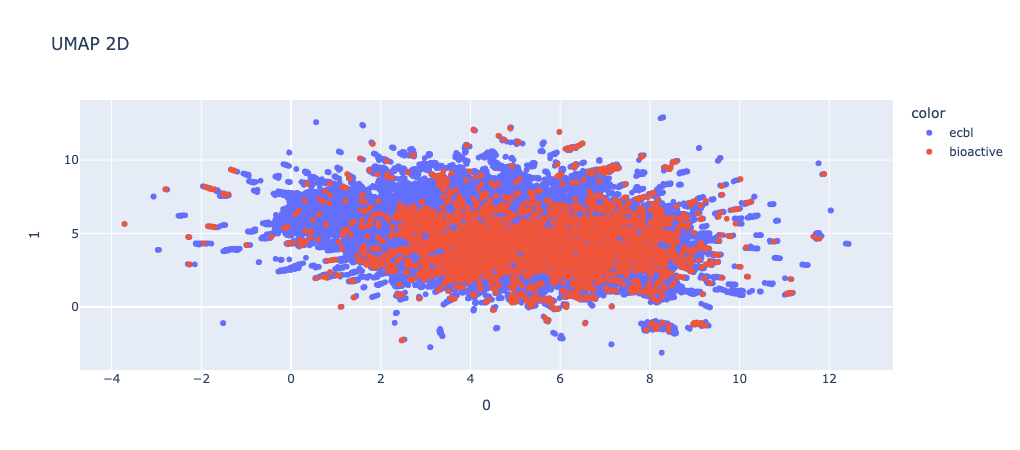

**Figure S5: Feature importance of MHFP6 fingerprints.** The rows represent the MHFP6 bits, and the columns represent the five label classes for the model. The colours in the heatmap denote the importance of the feature in the respective label class.

**Tables**

| **Pathogen class** | **Activity** | **# Compounds with Ro5 violations (%)** | **# Maximum structural alerts** | **# structural alerts**  **(%)** | **# No structural alerts (%)** |
| --- | --- | --- | --- | --- | --- |
| Gram-positive | Active | 11511 | 9 | 11,783 (67.22) | 5,747 (32.78) |
|  | Inactive | 21450 | 8 | 20,419 (71.48) | 8,149 (28.52) |
| Gram-negative | Active | 8651 | 9 | 7,764 (67.03) | 3,819 (32.97) |
|  | Inactive | 22146 | 12 | 21,158 (71.93) | 8,257 (28.07) |
| Acid-fast | Active | 3,958 | 7 | 2,741 (69.25) | 1,217 (30.75) |
|  | Inactive | 7,631 | 8 | 5,219 (68.39) | 2,412 (31.61) |
| Fungi | Active | 3288 | 8 | 2,989 (69.80) | 1,293 (30.20) |
|  | Inactive | 11492 | 8 | 9,581 (69.52) | 4,201 (30.48) |

**Table S1: Summary of Ro5 violations and structural alerts in AMR-KG.** For each pathogen class, the compounds have been distributed into active and inactive classes based on the pChEMBL threshold of 5.

| **Fingerprint** | **Model** | **Kappa score (Classic trained)** | **Kappa score (SMOTE trained)** |
| --- | --- | --- | --- |
| ECFP8 | Random forest | **0.62** | **0.81** |
|  | XGBoost | 0.58 | 0.72 |
|  | Light GBM | 0.54 | 0.67 |
|  | Decision Tree | 0.46 | 0.70 |
|  | Logistic Regression | 0.41 | 0.51 |
|  | Naive Bayes | 0.27 | 0.31 |
| RDKit | Random forest | **0.62** | **0.83** |
|  | XGBoost | 0.62 | 0.78 |
|  | Light GBM | 0.56 | 0.70 |
|  | Decision Tree | 0.48 | 0.74 |
|  | Logistic Regression | 0.41 | 0.51 |
|  | Naive Bayes | 0.17 | 0.17 |
| MACCS | Random forest | **0.61** | **0.81** |
|  | XGBoost | 0.56 | 0.68 |
|  | Decision Tree | 0.52 | 0.73 |
|  | Light GBM | 0.51 | 0.62 |
|  | Logistic Regression | 0.29 | 0.34 |
|  | Naive Bayes | 0.10 | 0.14 |
| MHFP6 | XGBoost | **0.62** | 0.80 |
|  | Random forest | 0.62 | **0.84** |
|  | Light GBM | 0.60 | 0.75 |
|  | Decision Tree | 0.44 | 0.67 |
|  | Logistic Regression | 0.42 | 0.57 |
|  | Naive Bayes | 0.14 | 0.16 |
| ErG | Random forest | **0.60** | **0.81** |
|  | XGBoost | 0.56 | 0.70 |
|  | Decision Tree | 0.51 | 0.70 |
|  | Light GBM | 0.52 | 0.66 |
|  | Logistic Regression | 0.29 | 0.34 |
|  | Naive Bayes | 0.06 | 0.08 |
| ChemPhys | Random forest | **0.54** | **0.72** |
|  | XGBoost | 0.47 | 0.58 |
|  | Decision Tree | 0.42 | 0.57 |
|  | Light GBM | 0.44 | 0.52 |
|  | Logistic Regression | 0.13 | 0.22 |
|  | Naive Bayes | 0.06 | 0.07 |

**Table S2: Evaluation metrics on hold-out set for all chemical fingerprints across six ML models, namely XGBoost, random forest, decision tree, logistic regression, light GBM and Naive Bayes.** The training set used for the models was generated using both “with” and “without” the SMOTE strategy. In bold are the top metrics for the fingerprint-training method combination across the six models.

| **Model name** | **Hyperparameters** |
| --- | --- |
| Random Forest | Num estimators = 100 to 500 with a step size of 100  Max depth = 10 to 110 with a step size of 20  Min sample split = 2 to 10 with a step size of 2  Min sample leaf = 1 to 4 |
| XGBoost | Num estimators = 1000  ETA = 0.01 to 0.1  Max dept = 2 to 10  Colsample by tree = 0.1 to 0.7 |

**Table S3: Hyperparameters used for optimizing the top two models: Random forest and XGBoost.**

| **Descriptors** | **Accuracy** | **Cohen Kappa** | **Macro precision** | **Macro recall** | **Macro F1** | **AUC-ROC** |
| --- | --- | --- | --- | --- | --- | --- |
| RDKIT | 0.743 | 0.658 | 0.751 | 0.744 | 0.748 | 0.837 |
| MHFP6 | 0.732 | 0.645 | 0.736 | 0.736 | 0.734 | 0.832 |
| MACCS | 0.706 | 0.614 | 0.692 | 0.719 | 0.703 | 0.821 |
| ECFP8 | 0.686 | 0.587 | 0.680 | 0.691 | 0.682 | 0.804 |
| ErG | 0.658 | 0.553 | 0.642 | 0.671 | 0.651 | 0.791 |
| ChemPhys | 0.634 | 0.519 | 0.610 | 0.631 | 0.617 | 0.768 |

**Table S4: XGBoost model performance on the test data.** The results shown in the table are the average metric reported across all four bacterial strain classes

| **Label Class** | **TPR** | **TNR** | **PPV** | **NPV** | **ACC** |
| --- | --- | --- | --- | --- | --- |
| Acid-fast | 0.814 | 0.985 | 0.852 | 0.980 | 0.968 |
| Fungi | 0.781 | 0.972 | 0.804 | 0.968 | 0.947 |
| Gram-negative | 0.653 | 0.963 | 0.762 | 0.939 | 0.916 |
| Gram-positive | 0.796 | 0.895 | 0.807 | 0.888 | 0.860 |
| Inactive | 0.737 | 0.859 | 0.652 | 0.901 | 0.827 |

**Table S5**: **MHFP6-RF test metrics.** The metrics reported in the table are as follows: true positive rate (TPR), true negative rate (TNR), positive predictive value or precision (PPV), negative predictive value (NPV), and accuracy (ACC).

| **Label Class** | **ECBL Model HitRate**  **(Experimental HitRate)** | **Bioactive Model HitRate**  **(Experimental HitRate)** |
| --- | --- | --- |
| Fungi | 1.424% (1.128%) | 45.918% (6.434%) |
| Gram-negative | 0.315% (0.028%) | 10.476% (1.055%) |
| Gram-positive | 1.096% (1.027%) | 15.000% (6.454%) |

**Table S6**: **EU-OS library prediction for each pathogen class.** For each pathogen class, the HitRate of the model and experiment are reported.

| **Enamine Catalog ID** | **Structure** | **Predicted Label** | **Probability** | **Cosine similarity** |
| --- | --- | --- | --- | --- |
| Z2583154272 | 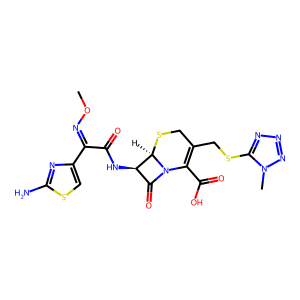 | Gram-negative | 0.876379 | 0.90 |
| Z1741977144 | 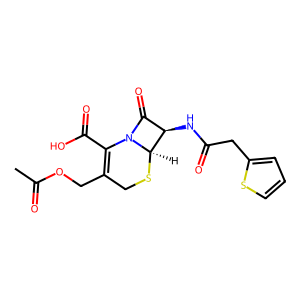 | Gram-positive | 0.838351 | 1.00 |
| Z1143251035 | 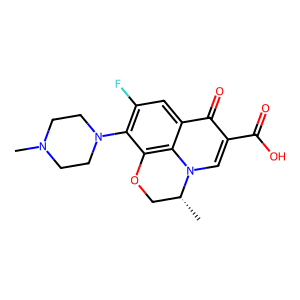 | Gram-positive | 0.807160 | 1.00 |
| Z2944548679 | 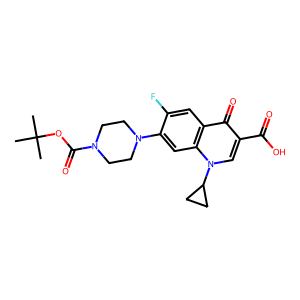 | Gram-positive | 0.804423 | 1.00 |
| Z1515385072 | 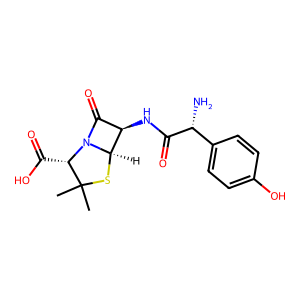 | Fungi | 0.753295 | 1.00 |
| Z2327029147 | 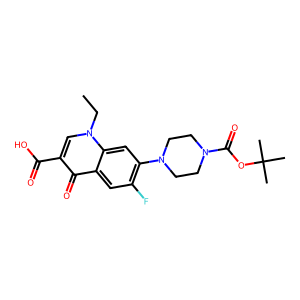 | Gram-positive | 0.634375 | 0.89 |
| Z1269203097 | 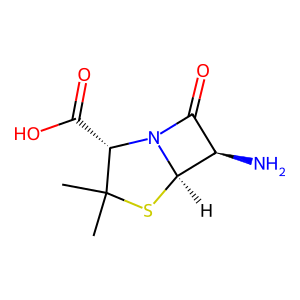 | Inactive | 0.593274 | 1.00 |
| Z647975500 | 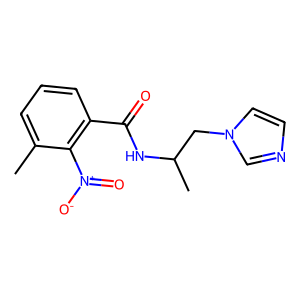 | Fungi | 0.586958 | 0.74 |
| Z3016338143 | 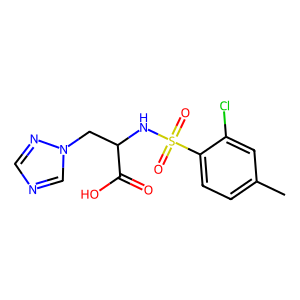 | Fungi | 0.572667 | 0.72 |
| Z27988606 | 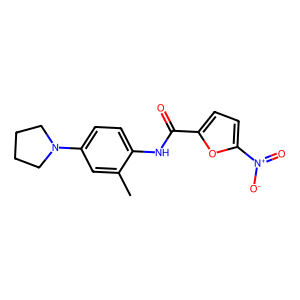 | Acid-fast bacteria | 0.559644 | 0.79 |
| Z1665472861 | 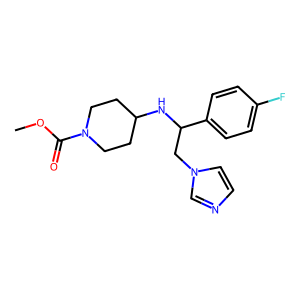 | Fungi | 0.556708 | 0.74 |
| Z27280112 | 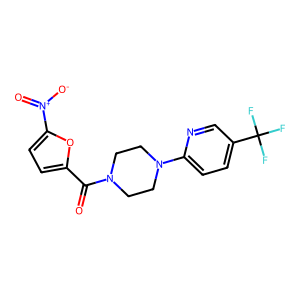 | Acid-fast bacteria | 0.548675 | 0.82 |
| Z85910719 | 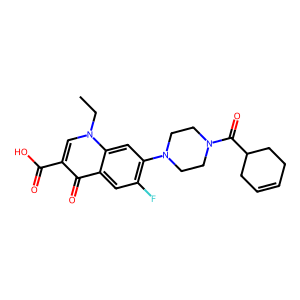 | Gram-positive | 0.542699 | 0.86 |
| Z85910185 | 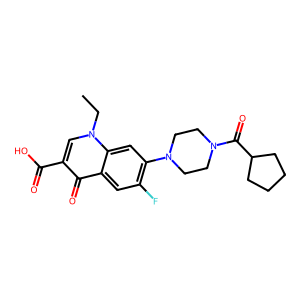 | Gram-positive | 0.542338 | 0.88 |
| Z1023434358 | 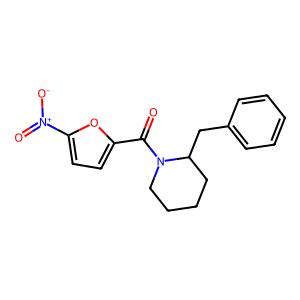 | Acid-fast bacteria | 0.534643 | 0.75 |
| Z969651104 | 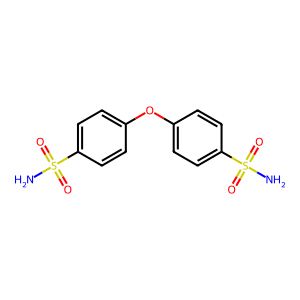 | Inactive | 0.533619 | 0.81 |
| Z1127188754 | 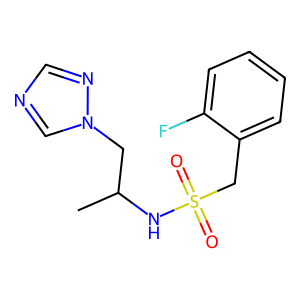 | Fungi | 0.528839 | 0.72 |
| Z1159338421 | 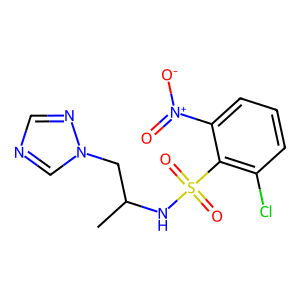 | Fungi | 0.525438 | 0.72 |
| Z1171969599 | 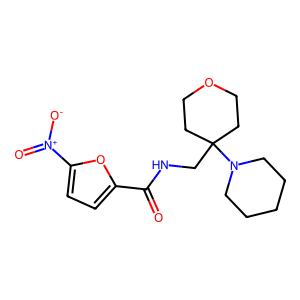 | Acid-fast bacteria | 0.523021 | 0.78 |
| Z85910290 | 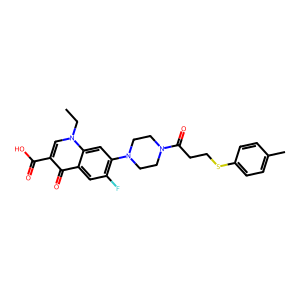 | Gram-positive | 0.522571 | 0.84 |
| Z970066380 | 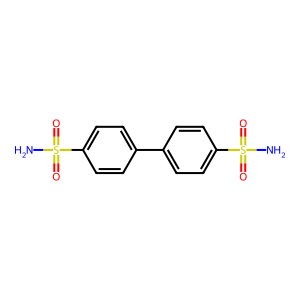 | Inactive | 0.522452 | 0.82 |
| Z1159338558 | 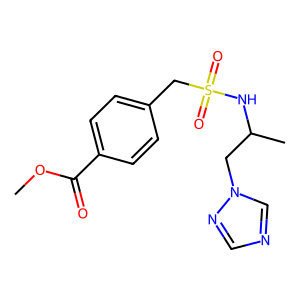 | Fungi | 0.522333 | 0.72 |
| Z137391786 | 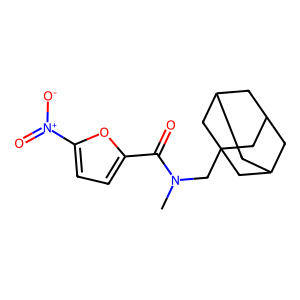 | Acid-fast bacteria | 0.519121 | 0.75 |
| Z57704153 | 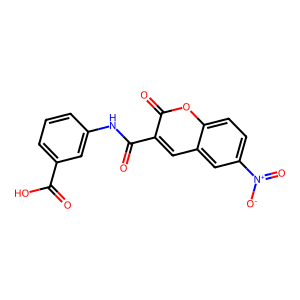 | Inactive | 0.517500 | 0.93 |
| Z85910288 | 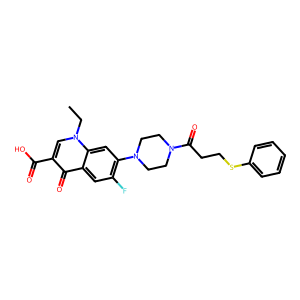 | Gram-positive | 0.516940 | 0.85 |
| Z414626052 | 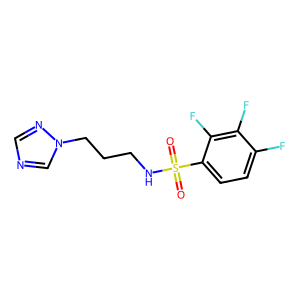 | Fungi | 0.515613 | 0.71 |
| Z1568296378 | 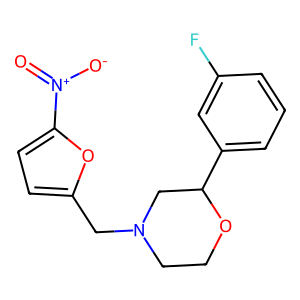 | Acid-fast bacteria | 0.514833 | 0.82 |
| Z799986482 | 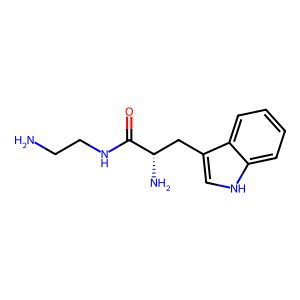 | Inactive | 0.513149 | 0.78 |
| Z414649268 | 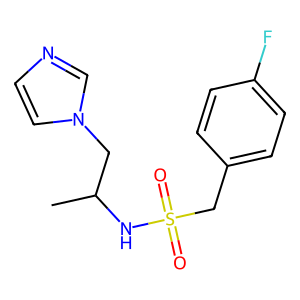 | Fungi | 0.505649 | 0.72 |
| Z416345606 | 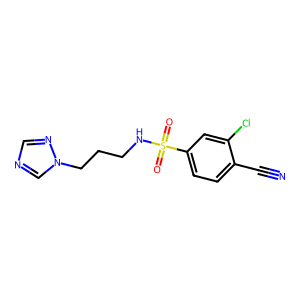 | Fungi | 0.504167 | 0.72 |
| Z1083844456 | 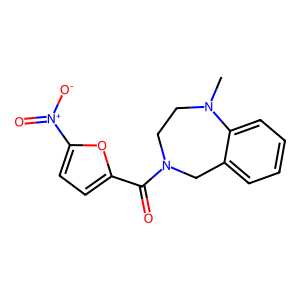 | Acid-fast bacteria | 0.503458 | 0.76 |
| Z2435971107 | 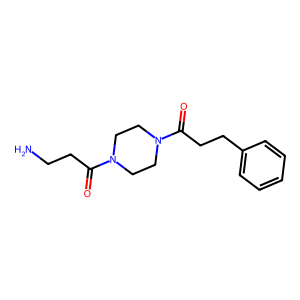 | Inactive | 0.501583 | 0.74 |
| Z1127188777 | 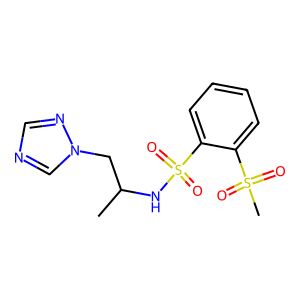 | Fungi | 0.500062 | 0.71 |

**Table S7**: **High confidence prediction of Enamine library.** The high-confidence predictions here are considered to have a prediction threshold greater than 0.5. We also demonstrate the MHFP6 fingerprint cosine similarity of these Enamine compounds with the training dataset.
